# Disruption of PI3K-OxPhos Coupling by Trehalose Drives a BCAA-to-Lipid Metabolic Switch in Hormone-Receptor–Positive Breast Cancer

**DOI:** 10.64898/2026.04.14.718262

**Authors:** Tashvinder Singh, Prabhsimran Kaur, Anjana Munshi, Sandeep Singh

## Abstract

Cancer cells sustain proliferation through dynamic coordination between mitochondrial oxidative phosphorylation (OxPhos) and anabolic carbon metabolism. How this metabolic coupling can be selectively destabilized in subtype-specific contexts remains poorly defined. Here we identify trehalose, a disaccharide previously linked to autophagy modulation, as a regulator of mitochondrial–anabolic integration in breast cancer. Using high-resolution respirometry, untargeted metabolomics, and signalling analyses across estrogen/progesterone receptor–positive (ER⁺) and triple-negative models, we show that trehalose preferentially impairs mitochondrial bioenergetics in OxPhos-dependent ER⁺ cells. Trehalose reduced electron transport system capacity, NADH-linked respiration, mitochondrial membrane potential, and coupling efficiency, while suppressing mitochondrial biogenesis markers. These bioenergetic effects coincided with attenuation of PI3K/Akt signalling and induction of p21-associated growth arrest. Metabolomic profiling revealed a coordinated redistribution of carbon flux characterized by depletion of branched-chain amino acids (BCAAs) and glycolytic intermediates alongside accumulation of long-chain fatty acids and cholesterol. Correlation network analysis uncovered a strong inverse relationship between BCAA-linked metabolism and lipid abundance, indicating a regulated metabolic trade-off rather than nonspecific stress. Functionally, trehalose enhanced the efficacy of mitochondrial-interfering agents such as tamoxifen and colchicine, while exerting minimal effects in metabolically flexible triple-negative cells. Together, these findings define trehalose as a metabolic modulator that constrains mitochondrial plasticity and enforces a lipid-buffered, growth-restrictive state in ER⁺ breast cancer, revealing a therapeutic vulnerability linked to mitochondrial dependency.

## Introduction

Breast cancer progression is driven by extensive metabolic reprogramming that enables tumour cells to adapt to fluctuating nutrient availability, hypoxia, and therapeutic stress. Central to this metabolic plasticity is mitochondrial bioenergetics, which integrates governs ATP production, redox balance, biosynthetic precursor generation with growth-promoting signaling networks [1, 2, 3, 4, 5]. The remarkable observation made by Warburg, ‘respiration damage’, emerged as a critical player in tumorigenesis [6]. However, rather than existing in a purely glycolytic state, breast cancer cells operate along a continuum between glycolysis and oxidative phosphorylation (OxPhos), with metabolic states varying across molecular subtypes and disease stages. This bioenergetic flexibility contributes to tumour heterogeneity, therapeutic resistance, and survival under metabolic stress [7, 8, 9, 10, 11, 12, 13, 14]. Mitochondrial dependency is particularly pronounced in estrogen receptor–positive (ER⁺) breast cancers, which frequently retain robust OxPhos capacity and mitochondrial integrity, in contrast to more metabolically flexible triple-negative subtypes. Increasing evidence indicates that mitochondrial function is not merely supportive but acts as a regulatory hub coordinating anabolic metabolism, redox homeostasis, and stress adaptation. Disrupting this mitochondrial–anabolic coupling therefore represents a promising, yet underexplored, therapeutic strategy [15, 16].

Our previous work and others have shown that both genetic and post-transcriptional mechanisms can reprogram mitochondrial metabolism in breast cancer, altering respiratory efficiency and influencing therapeutic sensitivity. However, pharmacological modulators that reshape mitochondrial network organization without directly targeting respiratory complexes remain poorly defined [17]. Trehalose, a naturally occurring non-reducing disaccharide, is widely recognised for its cytoprotective and autophagy-inducing properties [18, 19, 20, 21, 22]. Emerging evidence suggests that trehalose can influence mitochondrial homeostasis, oxidative stress, and protein turnover, yet its impact on mitochondrial bioenergetics and metabolic network architecture in cancer cells remains unclear [19, 23].

Here, we investigate whether trehalose can destabilize mitochondrial–anabolic integration in breast cancer. By combining high-resolution respirometry, signalling analysis, and untargeted metabolomics, we define how trehalose remodel mitochondrial function and carbon flux distribution across metabolically distinct breast cancer subtypes, revealing a subtype-specific metabolic vulnerability linked to mitochondrial dependency.

## Results

### Trehalose displays selective and context-dependent drug synergy in ER⁺ and TNBC

Trehalose suppressed proliferation across cancer cell types in a dose-dependent manner and reduced viability in both ER/PR⁺ MCF7 and TNBC MDA-MB-231 cells with IC₅₀ values <10 mM **(Fig. S1)**. To test whether trehalose modifies therapeutic sensitivity, we quantified drug–drug interactions using Compusyn and Combenefit (HSA framework) across mechanistically diverse agents (**Fig. S2-S5**). A detailed synergism and antagonism analysis was performed in both MCF7 cells and MDA-MB-231 cells. Synergy was defined by convergent metrics (positive synergy scores, positive log (Fa/Fu), and CI < 1), whereas negative scores and CI > 1 indicated antagonism (**Fig. S4-S5**).

Across both cell lines, trehalose showed robust synergy primarily with tamoxifen and colchicine, whereas combinations with doxorubicin, erlotinib, etoposide, and camptothecin were largely additive-to-antagonistic **(Fig. 1A-C)**. 2-DG exhibited moderate additive-to-synergistic behavior. Paclitaxel displayed subtype divergence, with synergy in MCF7 but antagonism in MDA-MB-231 (Fig. 1A). Overall, synergy metrics were more pronounced in MCF7 cells, suggesting a stronger dependency on the cellular processes perturbed by trehalose in ER⁺ contexts. This suggests that the drugs showing synergism are more effect in combination while antagonism were found to be more effective as single-agent doses. The therapeutic impact of the trehalose was found to be more potent against MCF7 cells (**Fig. 1D**). We further analysed the impact of Trehalose on cellular and physiological parameters associated with different hallmarks of ER^+ve^/PR^+ve^ BC.

**Figure 1:**
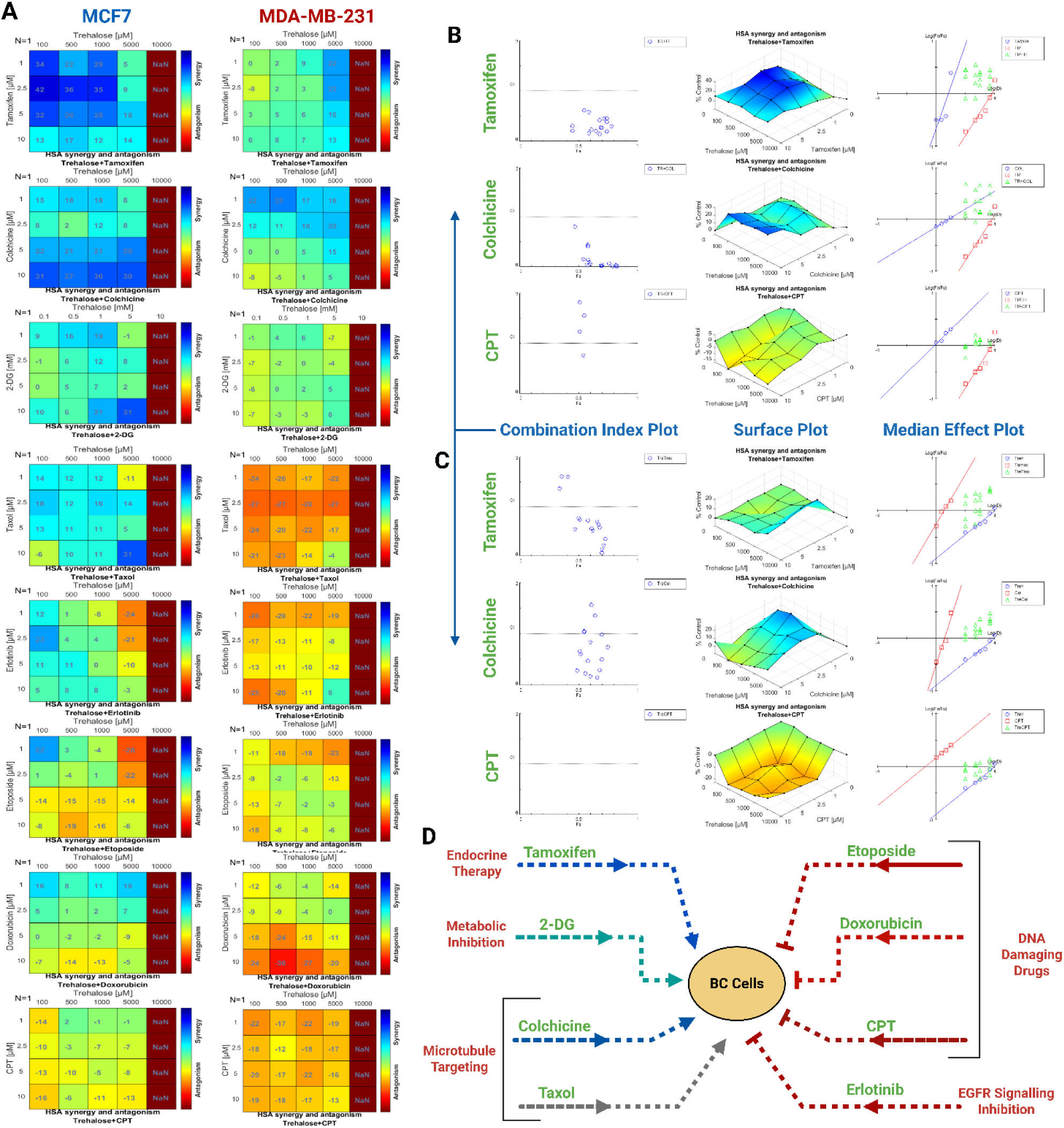
The synergistic or antagonistic cytotoxic impact of trehalose in combination with different drugs using HSA (Highest Single Agent) based Combenefit and Compusyn software analysis. **(A)** The higher blue colour intensity (positive value) indicates more synergistic combinations, while higher red colour (negative value) indicates high antagonistic combinations; intermediate values determine additive effects, while values near zero indicate no cytotoxic effects for the given combination. **(B)** The synergistic or antagonistic cytotoxic impact of trehalose in combination with different drugs on MCF7 cells using HSA (Highest Single Agent) based compusyn software analysis. The combination index plot, surface plot and median effect plot for trehalose in combination with different drugs. The combination index plotted CI values close to zero (near the x-axis) and less than 1, which signifies the combination to be synergistic, while more than 2 signifies the antagonistic effects for given combinations. **(C)** The synergistic or antagonistic cytotoxic impact of trehalose in combination with different drugs on MDA-MB-231 cells using similar HSA (Highest Single Agent) method. **(D)** The schematic diagram showing comprehensive synergism/antagonism of Trehalose combination with different molecular anti-cancer therapies. Blue dotted lines indicate synergism, grey dotted line indicate BC-subtype context dependent synergism/antagonism, while red line indicate antagonism against both ER/PR^=ve^ and TNBC.

### Trehalose disrupts mitochondrial physiology and imposes a stress-associated cell-cycle checkpoint in MCF7 cells

Trehalose reduced ROS levels in MCF7 cells in a dose-dependent manner (**Fig. 2A**) and lowered mitochondrial membrane potential (MMP), consistent with mitochondrial depolarization (**Fig. 2B**). Trehalose is known to induce autophagy and necrosis, because MMP and ROS are direct readouts of electron transport activity and redox homeostasis, these findings suggested impaired mitochondrial function [24, 25, 26]. Moreover, MMP and ROS production are crucial indicators of normal mitochondrial functioning; therefore, we analysed the impact of Trehalose on mitochondrial membrane integrity. To evaluate mitochondrial membrane integrity, we performed high-resolution respirometry with cytochrome c induction [27, 28].

**Figure 2:**
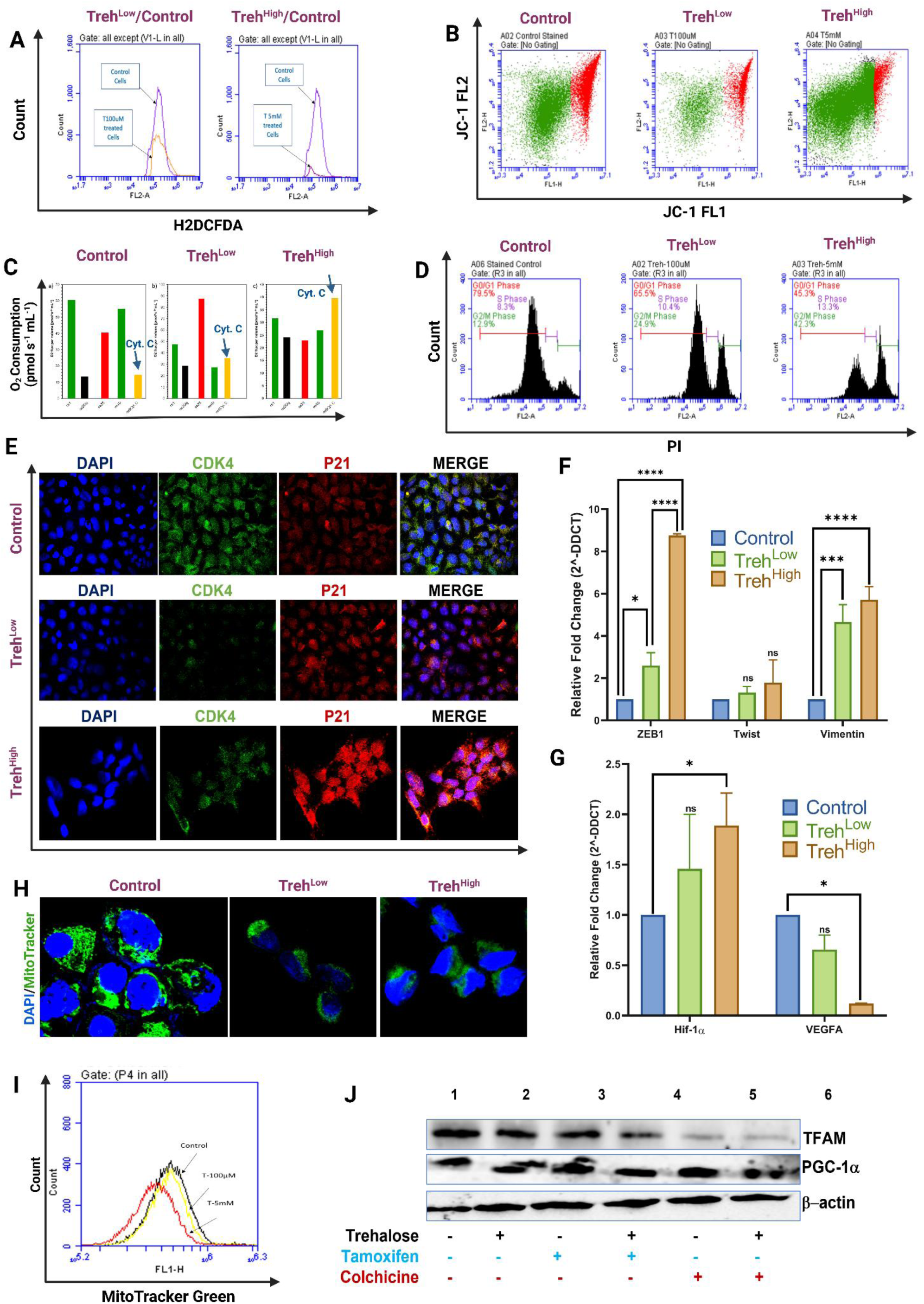
Trehalose disrupts the protective physiological functioning of ER/PR positive breast cancer cells. **(A)** The impact of trehalose on ROS production in MCF7 cells in a dose-dependent manner. **(B)** Trehalose depolarised mitochondrial membrane potential in a dose-dependent manner. **(C)** Trehalose induces an increase in basal OCR upon addition of cytochrome C, indicating a loose inner mitochondrial membrane in MCF7 cells. **(D)** Trehalose reduced cell cycle progression by arresting the cell cycle at G2/M stage. **(E)** Trehalose reduced the expression of CDK4, while increasing the expression of p21, a protein involved in the cell cycle checkpoint at the G2/M stage. **(F)** Trehalose induced the expression of ZEB1, Twist, and Vimentin, genes responsible for the stemness property and metabolic senescence. **(G)** Trehalose reduced the expression of VEGFA while inducing mRNA expression of Hif-1α. **(H)** Immunofluorescence analysis shows trehalose reduced mitochondrial count in MCF7 cells in a dose-dependent manner. **(I)** Flow cytometric analysis shows trehalose reduced mitochondrial number in MCF7 cells in a dose-dependent manner. The percentage of cell population stained for Mitotracker green was 54.78%, 34.94% in control and Treh^High^ MCF7 cells, respectively. **(J)** The impact of trehalose in combination with tamoxifen and colchicine on the protein expression of TFAM and PGC-1α.

After achieving the routing respiration, Cyt. C was added to evaluate change in OCR. Cytochrome c increased oxygen flux in MCF7^Treh^ cells in dose-dependent manner (**Fig. 2C**), indicating compromised outer mitochondrial membrane integrity and impaired respiratory compartmentalization. This loss of membrane integrity aligned with the observed reductions in MMP and ROS and supported a direct inhibitory effect of trehalose on mitochondrial function.

In addition, the effects of trehalose on key physiological parameters linked to tumorigenic potential and mitochondrial function were evaluated in MCF7 cells. Specifically, the proportion of cells in the G0/G1 phase decreased from 79.5% in control cells to 65.5% and 45.3% following Treh^High^ and Treh^Low^ treatments, respectively. Conversely, the S-phase population increased from 8.3% in control cells to 10.4% and 13.3% in Treh^High^ and Treh^Low^ cells, respectively. A pronounced accumulation of cells in the G2/M phase was also observed, rising from 12.9% in control cells to 24.9% and 42.3% in Treh^High^ and Treh^Low^ treatment groups, respectively (**Fig. 2D**). Trehalose altered cell-cycle progression, reducing the G0/G1 fraction and increasing S and G2/M populations, consistent with checkpoint activation under bioenergetic stress. This shift was accompanied by decreased CDK4 in dose-dependent and induction of p21 at high dose (**Fig. 2E**), linking mitochondrial dysfunction to growth arrest in MCF7 cells.

Interestingly, trehalose increased expression of EMT/stemness-associated markers (ZEB1, Twist, Vimentin) in MCF7 cells (**Fig. 2F**). However, hypoxia-linked transcriptional changes suggested a stress-adaptive program rather than a pro-angiogenic switch: HIF-1α increased, while VEGFA decreased (**Fig. 2G**). This pattern is compatible with a pseudo-hypoxic stress response coupled to growth restriction. These stemness markers are associated with increased metastasis as well as hypoxia/Hif-1α-driven EMT [29, 30, 31].

This suggests a trehalose-induced hypoxic condition correlates with low oxygen availability along with metabolic senescence-like phenotype and a reduction in ROS production [32, 33, 34], which also might be linked to reduced OxPhos in MCF7^Treh^ cells. Further, the impact of trehalose on mitochondrial functioning and OxPhos was assessed to gain wholistic mechanism.

### Trehalose suppresses mitochondrial abundance/biogenesis and triggers mitochondria-dependent cytotoxicity in ER⁺ cells

We next examined how Trehalose influences mitochondrial function. Mitotracker-based flow cytometry and confocal imaging demonstrated a dose-dependent reduction in mitochondrial staining intensity and mitochondrial content in trehalose-treated MCF7 cells (**Fig. 2H–I**). Consistent with reduced mitochondrial maintenance capacity, TFAM expression decreased with trehalose and was further reduced when trehalose was combined with tamoxifen or colchicine (**Fig. 2J**). PGC-1α remained largely unchanged with trehalose monotherapy and decreased primarily under combination conditions (**Fig. 2J**), suggesting that trehalose might preferentially disrupts mitochondrial maintenance/mtDNA specific ETC machinery and that combinatorial stress amplifies broader biogenesis suppression.

Our unpublished findings have already demonstrated the mitochondrial-dependent and independent cytotoxicity of tamoxifen and colchicine in MCF7 cells respectively. This further validates the fact that Trehalose induced synergistic inhibition of mitochondrial biogenesis when given in combination with tamoxifen and colchicine. Further, we analysed the impact of trehalose on the mechanistic pathways associated with mitochondrial functioning and metabolic plasticity of MCF7 cells. Trehalose activated apoptotic machinery, with increased caspase processing (caspase-9, -7, -3) that was enhanced by combination with tamoxifen or colchicine (Fig. 3A). High-dose trehalose also induced a modest necrotic fraction (Fig. 3B), consistent with severe bioenergetic stress producing mixed death modalities.

**Figure 3:**
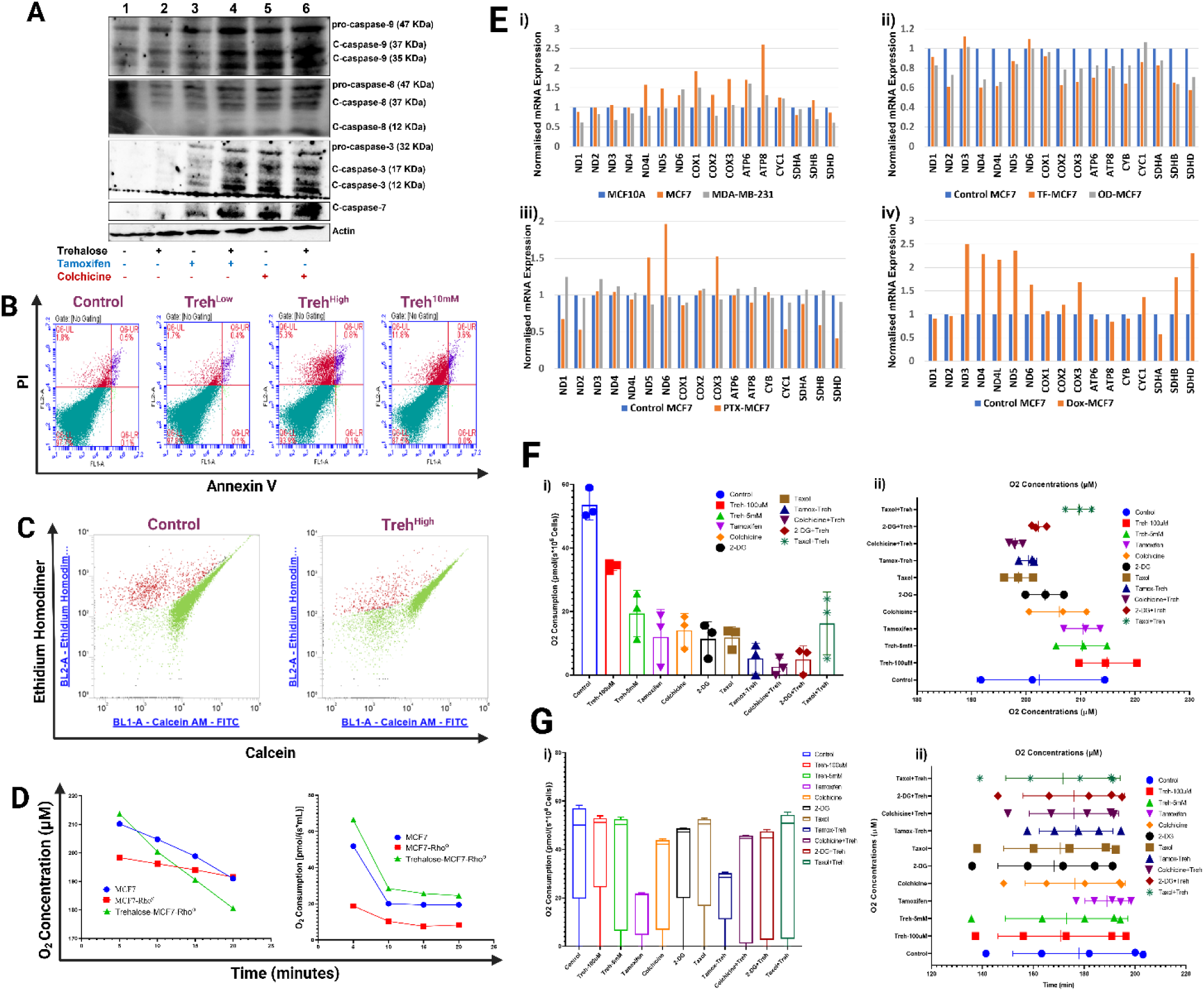
Trehalose inhibited mitochondrial biogenesis and mediates mitochondrial-dependent anti-cancer effects in MCF7 cells. Further, Trehalose reduced elevated mitochondrial respiration in ER/PR positive breast cancer cells. **(A)** Trehalose induced the protein expression of caspase-3, caspase-7, caspase-8, and caspase-9 involved in apoptotic machinery in MCF7 cells. **(B)** The flow cytometry analysis of the Pi/Annexin assay shows trehalose-induced cell death in MCF7 cells at higher concentrations. **(C)** The flow cytometry analysis of the Live/Dead cell assay shows that trehalose had no cytotoxic effect on MCF7-ρ0 cells. **(D)** Oroboros-based O2k Fluo-respirometry analysis shows no difference in basal OCR upon treatment with trehalose in MCF7-ρ0 cells. **(E)** Reduced expression of mitochondrial encoded genes in BC subtypes and their response to anti-cancer therapy. The transcriptomic counts of mitochondrial encoded genes in i) MCF, MDA-MB-231, and MCF10A cells; ii) tamoxifen-treated MCF7 cells; iii) taxol-treated MCF7 cells; iv) doxorubicin-treated MCF7 cells. **(F)** High resolution respirometry shows a change in i) basal oxygen consumption rate (OCR in pmol/s/mL) and ii) O concentration (µM) in MCF7 cells in response to mono and combinatorial therapy. **(G)** High resolution respirometry shows a change in i) basal oxygen consumption rate (OCR in pmol/s/mL) and ii) O concentration (µM) in MDA-MB-231 cells in response to mono and combinatorial therapy.

To directly test mitochondrial dependency, we used mtDNA-depleted MCF7 ρ⁰ cells. As ρ0 cells are resistant to apoptosis or mitochondrial-dependent cell death [35]. Trehalose did not significantly reduce viability in ρ⁰ cells (**Fig. 3C**) and had minimal impact on oxygen consumption in ρ⁰ cells relative to untreated ρ⁰ controls (**Fig. 3D**), indicating that trehalose-induced cytotoxicity requires functional mitochondria and is not explained by non-mitochondrial respiration. Thus, signifying a very low effect of trehalose on non-mitochondrial respiration (**Fig. 3D**).

To better understand how trehalose induces mitochondrial dysfunction, its impact on modulating mitochondrial metabolic sub-domains in ER/PR^+ve^ BC cells was assessed. But before assessing OxPhos parameters, we evaluated the alteration of mitochondrial gene expression in BC cells using publicly available RNA-seq data.

### Transcriptomic and clinical analyses support subtype-specific mitochondrial programs and prognostic relevance

Public RNA-seq analyses revealed differential expression of mtDNA encoded genes among different BC subtypes, and further distinguishes the impact of the synergistic/antagonistic drugs on these genes (**Fig. 3E; S6**). Relative to non-tumorigenic MCF10A, respiratory/OxPhos-associated transcripts were elevated in ER⁺ MCF7 and modest increase in TNBC MDA-MB-231 cells (**Fig. 3E-i**), consistent with higher mitochondrial reliance in ER⁺ cells. This displays metabolic heterogeneity as OxPhos is dysregulated during BC progression, and it is differentially active among BC subtypes [36]. Drug-associated transcriptomic modulation further indicated that agents synergizing with trehalose tend to suppress mitochondrial respiratory gene expression, whereas non-synergistic agents showed limited or inconsistent modulation (**Fig. 3E; S6**), reinforcing a mechanistic link between trehalose synergy and mitochondrial pathway suppression. The RNA-seq data also revealed that doxorubicin and tamoxifen increased and reduced the expression of mitochondrial gene encoding transcripts respectively in MCF7 cells. While taxol displayed modest decrease in mtDNA genes (**Fig. 3Eii-iii**). The etoposide was found not to affect the expression of these transcripts in MCF7 cells (**Fig. 3E-iv**). Survival analysis showed that high expression of several respiratory-chain–linked genes (SDHA, SDHB, CYC1, COX4I1, COX4I2, ATP8A2) associated with poorer overall survival (log-rank P < 0.05; HR > 1), whereas ATP8A1 associated with improved survival and SDHD showed no significant association (**Fig. S7**). These results support the clinical relevance of mitochondrial respiratory programs as biomarkers and potential vulnerabilities.

We next assessed basal oxygen consumption in MCF7 and MDA-MB-231 cells under trehalose and synergistic drug combinations (tamoxifen, colchicine, paclitaxel, 2-DG; **Fig. S8**) to identify the differential metabolic dependency and OxPhos heterogeneity towards therapeutics.

### Trehalose selectively suppresses mitochondrial respiration and OxPhos capacity in ER⁺ BC cells

The doses for the trehalose-drug combinations were chosen on the basis of their highest synergistic effect at the respective dosage combination. (**Fig. S8**). The oxygen consumption rate (OCR) values in pmol/s/10^6^ cells were also taken at 10-minute intervals between 0 and 20 minutes. The recorded OCR values changed from 59 to 50 for untreated MCF7 cells, 35 to 34 for Treh^Low^, 11 to 25 for Treh^High^, 2 to 18.6 for tamoxifen, 8 to 18 for colchicine, 5 to 13 for 2-DG, and 7.9 to 13.9 for Taxol, respectively. For the combinatorial doses, the OCR values changed from 0 to 9 for Treh-TF, 0 to 5.49 for Treh-Col, 0 to 7.5 for Treh-2DG, and 5 to 23 for Treh-PTX (**Fig. 3F-i**). The O2 concentration was rapidly decreased in control and Treh^Low^ -treated both MCF7 cells due to a high OCR compared to a moderate decrease in other treatment groups, specifically in combinatorial doses (**Fig. 3F-ii**).

In MDA-MB-231 cells, the recorded OCR values changed from 0 to 50 for control cells, 0 to 51 for Treh^low^, 0 to 50 for Treh^high^, 0 to 21 for tamoxifen, 0 to 42 for colchicine, 0 to 48 for 2-DG, and 0 to 50.6 for Taxol. For the combinatorial doses, the OCR values changed from 0 to 29 for trehalose-tamoxifen, 0 to 44.57 for trehalose-colchicine, 0 to 44.8 for trehalose-2-DG, and 0 to 50.9 for trehalose-taxol combination-treated MDA-MB-231 cells (**Fig. 3G-i)**. Similarly, the impact on O2 concentration can be seen in MDA-MB-231 cells **(Fig. 3G-ii**).

In MCF7 cells, trehalose reduced basal respiration and further suppressed oxygen consumption when combined with tamoxifen or colchicine (**Fig. 3F**). Oxygen depletion kinetics were correspondingly slower, consistent with reduced respiratory flux (Fig. 3F-ii). In contrast, MDA-MB-231 cells exhibited minimal trehalose-driven suppression of basal respiration; only tamoxifen alone or with trehalose reduced OCR appreciably (**Fig. 3G**), consistent with lower mitochondrial dependency in TNBC.

This might be due to either the differential impact of trehalose or the differential metabolic plasticity of ER/PR^+ve^ and TNBC. To evaluate this, trehalose was further analysed for its impact on specific mitochondrial parameters and sub-domains involved in regulating the energy demands.

### Trehalose decreased coupling efficiency and mitochondrial respiratory complex activity via inhibiting PI3K/Akt signaling in MCF7 cells

The impact of Trehalose on OxPhos complexes, leak respiration, ETS capacity, and coupling efficiency was analysed in both MCF7 and MDA-MB-231 cells. Using SUIT protocols, trehalose in MCF7 cells reduced FCCP-stimulated ETS capacity while leaving oligomycin-insensitive leak respiration relatively unchanged (**Fig. 4A; S8**). The L/E ratio was found to be 0.76 and 0.90 in low and Treh^high^ of Trehalose-treated cells, respectively, as compared to 0.52 in untreated MCF7 cells. This increased the leak-to-ETS ratio (L/E) (**Fig. 4B**), indicating reduced respiratory efficiency due primarily to constrained maximal electron transport capacity. L/E ratio values range between 0 and 1; near 0 reflects tight coupling/high efficiency; higher L/E indicates greater uncoupling/inefficiency [37, 38].

**Figure 4:**
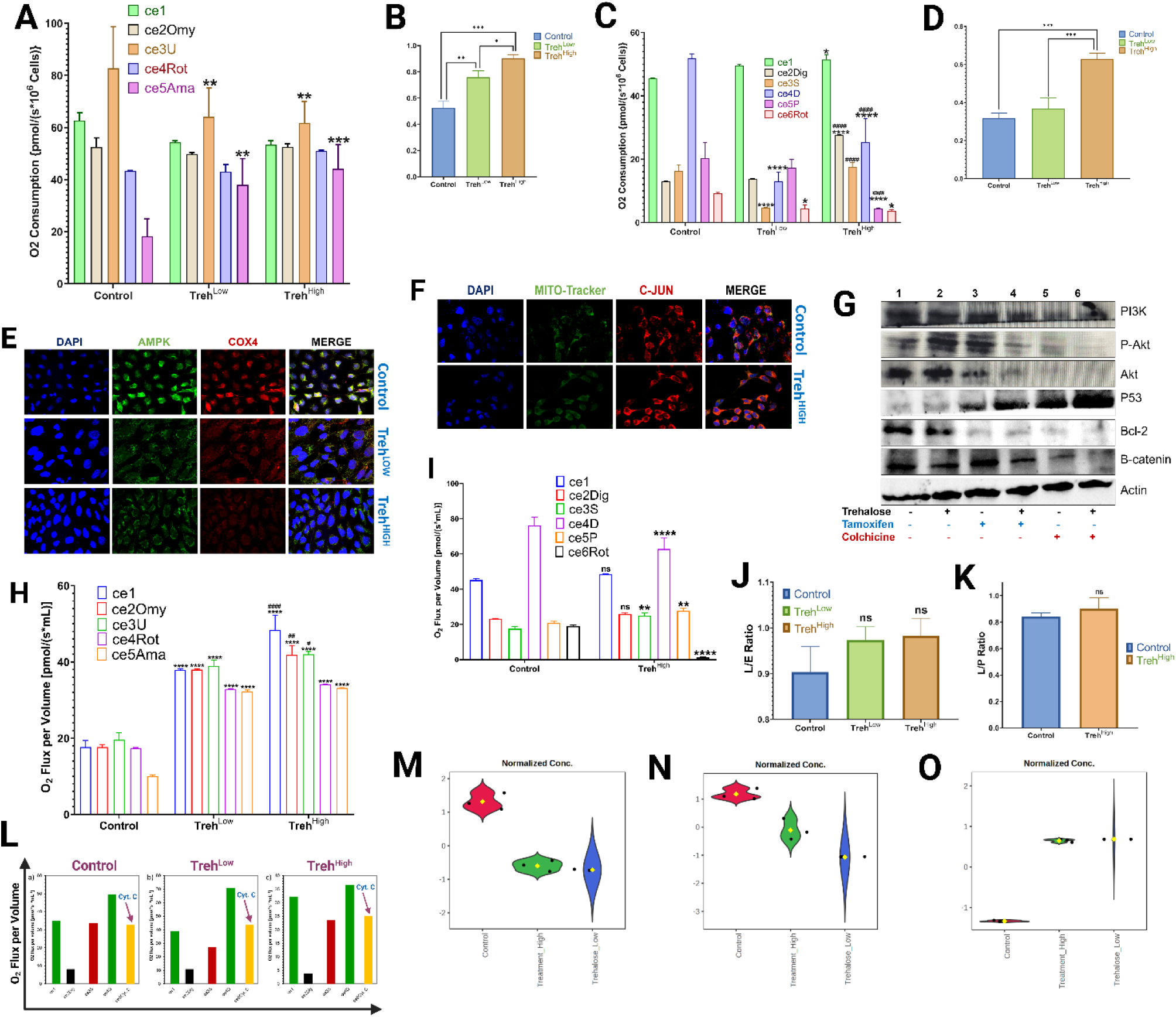
Trehalose reduced OxPhos by disrupting ETS capacity and NADH-linked respiration in MCF7 cells, while no significant inhibitory effect was observed in MDA-MB-231 cells. **(A)** The bar graph shows trehalose disrupts ETS-linked respiration by reducing maximal OxPhos capacity in MCF7 cells, as analysed by the addition of uncoupler FCCP. **(B)** The bar graph shows a reduction in the leak respiration to ETS ratio and induction of uncoupling in trehalose-treated MCF7 cells. **(C)** The bar graph shows trehalose disrupts NADH-linked respiration by reducing complex I activity in MCF7 cells, as analysed by the addition of pyruvate (complex I substrate). **(D)** The bar graph shows a reduction in the leak respiration to routine respiration ratio and induction of uncoupling in trehalose-treated MCF7 cells. **(E)** The immunofluorescence imaging shows that trehalose reduced the protein expression of AMPK and COX4 in MCF7 cells, part of metabolic homeostasis and OxPhos, respectively. **(F)** The immunofluorescence imaging shows that trehalose slightly increased the protein expression of c-Jun (metabolic sensor) in MCF7 cells. **(G)** The immunoblotting shows that trehalose alone and in combination with tamoxifen and colchicine reduced the protein expression of PI3K, Akt, p53, and β-catenin in MCF7 cells. **(H)** The bar graph shows trehalose does not have a significant impact on ETS-linked respiration in MDA-MB-231 cells, as analysed by the addition of the uncoupler FCCP. **(I)** The bar graph shows that trehalose does not have a significant impact on NADH-linked respiration in MDA-MB-231 cells, as analysed by the addition of pyruvate (a complex I substrate). **(J)** The bar graph shows no change in the leak respiration to ETS ratio in trehalose-treated MDA-MB-231 cells. **(K)** The bar graph shows no change in the leak respiration to routine respiration ratio in trehalose-treated MDA-MB-231 cells. **(L)** The bar graph shows no change in basal OCR after the addition of cytochrome C in trehalose-treated MDA-MB-231 cells. **(M)** Normalized metabolites levels across treatment conditions. Violin plot showing z-score–normalized lactic acid abundance in Control, Treatment_High, and Trehalose_Low groups. Points represent biological replicates and yellow diamonds indicate group means. **(N)** Violin plot showing z-score–normalized valine abundance across experimental conditions, highlighting relative changes in branched-chain amino acid levels. **(O)** Violin plot showing z-score–normalized cholesterol abundance across groups, indicating relative lipid enrichment under trehalose treatment.

The addition of digitonin, termed ROX state (residual oxygen consumption), and the O2 consumption was found to be increased in trehalose-treated cells in a dose-dependent manner. However, after adding ADP, the oxygen consumption was comparatively low in both Treh^High^ and Treh^Low^ treated MCF7 cells as compared to control cells. OCR was decreased in trehalose-treated cells in a dose-dependent manner after adding pyruvate (complex I substrate) and rotenone (complex I inhibitor) as compared to control cells (**Fig. 4C**). Trehalose had a low effect on inducing the change in leak respiration while downregulating pyruvate-induced NADH-linked respiration. Further, the impact of Trehalose on the activity of different mitochondrial respiratory complex was analysed in MCF7 cells. Complex I–linked respiration was particularly impaired, as shown by reduced NADH-linked flux after pyruvate stimulation and expected suppression with rotenone (**Fig. 4C**). Leak/Routine-associated indices were consistent with reduced oxidative efficiency (**Fig. 4D**). The L/P ratio was found to be 0.37 and 0.63 in Treh^High^ and Treh^Low^ Trehalose-treated cells, respectively, as compared to 0.32 in untreated MCF7 cells (**Fig. 4D**). We further analysed the expression of key metabolic signalling pathways responsible for regulating metabolic homeostasis.

Trehalose (Treh^Low^ and Treh^High^) also altered metabolic signalling and mitochondrial protein markers: AMPK and COX4 decreased, while c-Jun increased at higher trehalose (**Fig. 4E–F**), consistent with stress-adaptive remodeling. PI3K/Akt signalling (Akt and p-Akt) decreased, particularly under tamoxifen/colchicine ± trehalose, while p53 increased in the most growth-suppressive conditions (**Fig. 4G**). Collectively, these data support a model in which trehalose limits OxPhos capacity, constrains mitochondrial programme maintenance, and cooperates with mitochondrial-active therapeutics.

### Inhibitory effects of trehalose on coupling efficiency are no longer valid in MDA-MB-231 cells

Further, the leak respiration and ETS capacity both were found to be increased in MDA-MB-231^Treh^ cells (**Fig. 4H; S8**). However, in MDA-MB-231 cells, baseline respiration was comparatively highly uncoupled and trehalose produced only modest changes in ETS capacity and coupling indices (**Fig. 4H–K**). The L/E ratio was found to be 0.97 and 0.98 in Treh^Low^ and Treh^High^ MDA-MB-231 cells, respectively, as compared to 0.90 in untreated MDA-MB-231 cells (**Fig. 4I**). The OCR was significantly decreased and increased after the addition of ADP and pyruvate, respectively, and significantly reduced to zero after adding rotenone in trehalose-treated cells, as compared to control MDA-MB-231 cells (**Fig. 4J**). Similarly, the L/P ratio was calculated, and the L/P ratio was found to be 0.90 in Trehalose-treated cells as compared to 0.84 in untreated MDA-MB-231 cells (**Fig. 4K**).

Importantly, cytochrome c produced no significant OCR increase in Treh^High^ and Treh^Low^ MDA-MB-231 cells (**Fig. 4L**), indicating preserved mitochondrial membrane integrity and providing a functional basis for relative resistance to trehalose-driven respiratory suppression. It might be the reason for the low impact of trehalose on mitochondrial respiration in MDA-MB-231 cells compared to MCF7 cells. Similar to the effects of Trehalose in MCF7 cells, trehalose reduced the OxPhos capacity by limiting normal respiration and maximum efficiency of OxPhos, but to a lesser extent in MDA-MB-231 cells. Differential OxPhos capabilities of MCF7 and MDA-MB-231 cells establish the metabolic heterogeneity among BC subtypes, indicating metabolic-dependent therapeutic outcomes.

### Multivariate metabolomics identifies a BCAA/glycolysis/lipid axis as the dominant discriminator of trehalose response

Given the strong suppression of mitochondrial bioenergetics in MCF7 cells, we performed GC–MS metabolomics to define downstream metabolic adaptations. Z-score-normalized profiles showed a coordinated, dose-dependent shift (**Fig. 4M–O**): lactate decreased (reduced glycolytic output) (**Fig. 4M**), BCAAs (including valine) were depleted (**Fig. 4N**), and lipid-associated metabolites especially cholesterol and long-chain fatty acids accumulation (**Fig. 4O**). Hierarchical clustering separated control, TrehLow, and TrehHigh groups, indicating stable metabolic reprogramming rather than stochastic stress (**Fig. 5A**).

**Figure 5:**
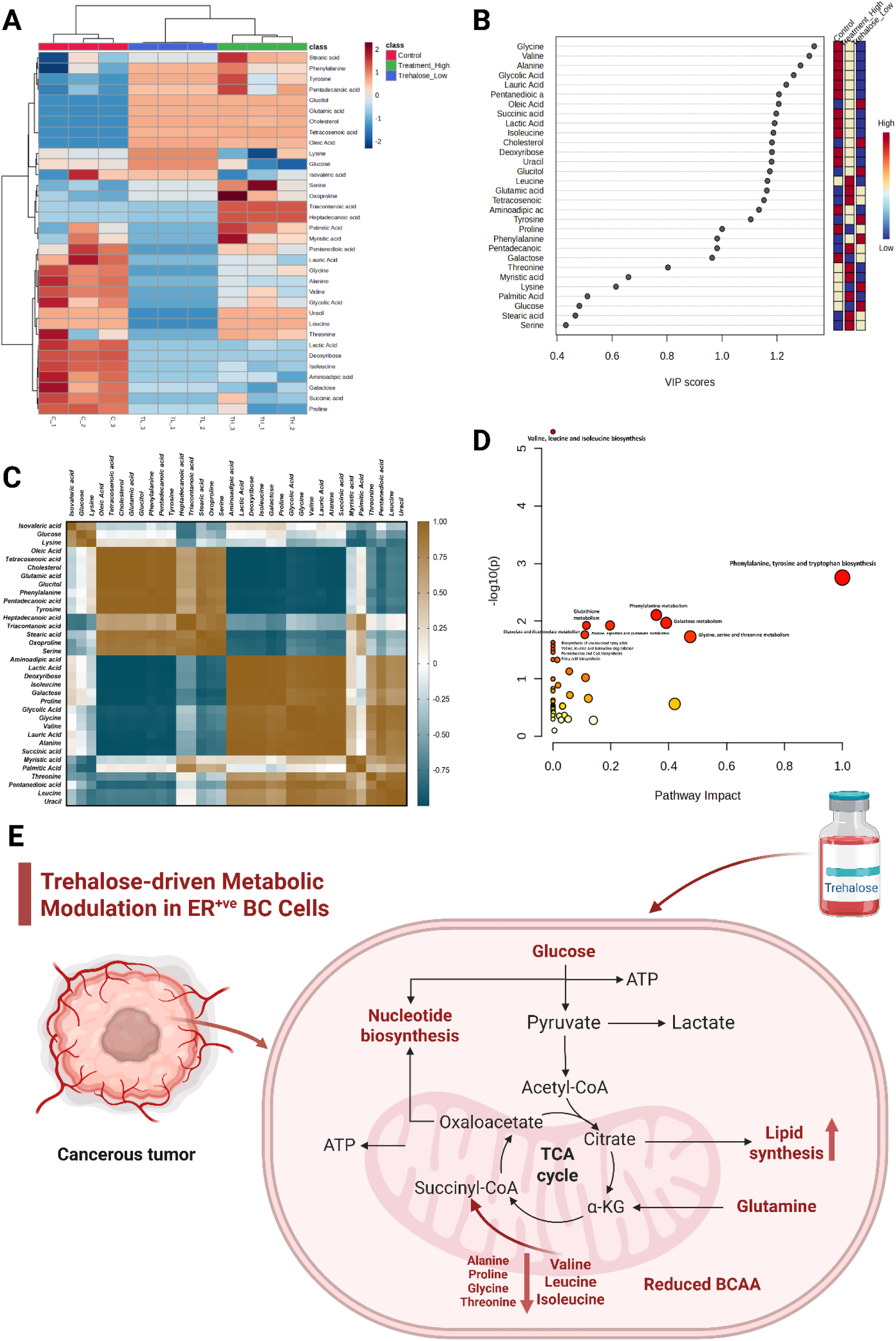
The metabolic vulnerabilities induced by Trehalose in MCF7 cells. **A)** Heatmap showing altered proportions of 32 metabolites in MCF7 cells under trehalose treatment in MCF7 cells. **B)** Correlation matrix showing positive and negative correlation among key altered significant metabolites under trehalose treatment. **C)** VIP scores for altered metabolites under therapeutic conditions and metabolic modulations. **D)** Significant pathways associated with altered metabolites in MCF7^Treh^ cells. **(E)** Schematic diagram showing impact of Trehalose on modulation metabolism in ER/PR^+ve^ BC cells.

The amino acids glycine, alanine, valine, leucine, isoleucine, proline, and threonine were significantly present in lower concentrations in trehalose-treated MCF7 cells, and their proportions decreased in dose dependent manner. On the other hand, amino acids such as phenylalanine, tyrosine, glutamic acid, serine, and oxoproline were significantly present in higher concentrations in trehalose-treated MCF7 cells. The lipids and long-chain fatty acids, including stearic acid, cholesterol, pentadecanoic acid, tetracosanoic acid, oleic acid, pentanedioic acid, palmitic acid, myristic acid, tricontanoic acid, and heptadecanoic acid, were significantly present in higher proportions in dose dependent manner in trehalose-treated MCF7 cells. Trehalose treatment suppressed glycolytic and branched-chain amino acid–associated metabolites while inducing accumulation of long-chain fatty acids and lipid-associated species, consistent with a shift from an OxPhos-supported proliferative state to a lipid-loaded, bioenergetically constrained phenotype (**Fig. 5A**).

PLS-DA VIP analysis (multivariate model using partial least squares discriminant analysis; VIP > 1.2) identified the most discriminatory metabolites spanning BCAAs/amino acids (e.g., glycine, valine, alanine, and isoleucine), glycolysis/TCA-linked metabolites (e.g., lactate, succinate), and lipid/fatty-acid species (e.g., glycolic acid, lauric acid, pentanedioic acid, oleic acid, and cholesterol) (**Fig. 5B**). This multi-domain signature indicates that trehalose-driven state transitions reflect coordinated rewiring of carbon and nitrogen metabolism coupled to lipid homeostasis.

Pearson correlation analysis revealed distinct metabolic modules and metabolites with extreme positive and negative significant correlations [|r| ≥ 0.95, FDR-adjusted p < 0.05] were considered for biological interpretation, high confidence association and network construction. The analysis revealed two dominant modules (**Fig. 5C**). BCAAs and glycolysis-linked metabolites formed a positively correlated cluster (tight coupling between amino acid availability and pyruvate-linked metabolism). Lipid metabolites formed a second tightly correlated module that exhibited strong inverse correlations with the BCAA module (r < −0.95 for core nodes), consistent with a regulated metabolic trade-off: lipid accumulation rises as BCAA/glycolytic metabolites decline.

A strong positive correlation between amino acids was observed among isoleucine, proline, valine, alanine, and glycine (Group 1; r ≈ 0.99), and further correlation with other metabolites such as lactic acid, deoxyribose, galactose, and succinic acid (Group 2; r > 0.95), indicating tight coupling between BCAA availability and pyruvate-derived metabolites. In contrast, lipid metabolites formed a separate, tightly co-regulated cluster. Oleic acid exhibited a near-perfect positive correlation with tetracosanoic acid, pentadecanoic acid, glucitol, and cholesterol (Group 3; r ≈ 0.99), as well as glutamate, phenylalanine, and tyrosine (r > 0.90) (**Fig. 5C**). While negative correlations were seen among amino acids clusters (r < −0.95; **Cluster 1:** glutamate, phenylalanine and tyrosine; **Cluster 2:** isoleucine, proline, glycine, valine and alanine.

Enrichment and pathway impact analysis converged on amino acid metabolism (including BCAA degradation and glycine/serine/threonine pathways), galactose metabolism, lipid biosynthesis, Warburg effect and partial engagement of mitochondrial-linked shuttles (**Fig. 5D; Fig. S9–S10; Tables 1–2**), supporting a model of constrained central carbon flux with compensatory lipid buffering. Pathway impact analysis highlighted BCAA metabolism as well as lipid biosynthesis affected networks, with high impact values driven by central nodes rather than peripheral metabolites.

**Table 1:**
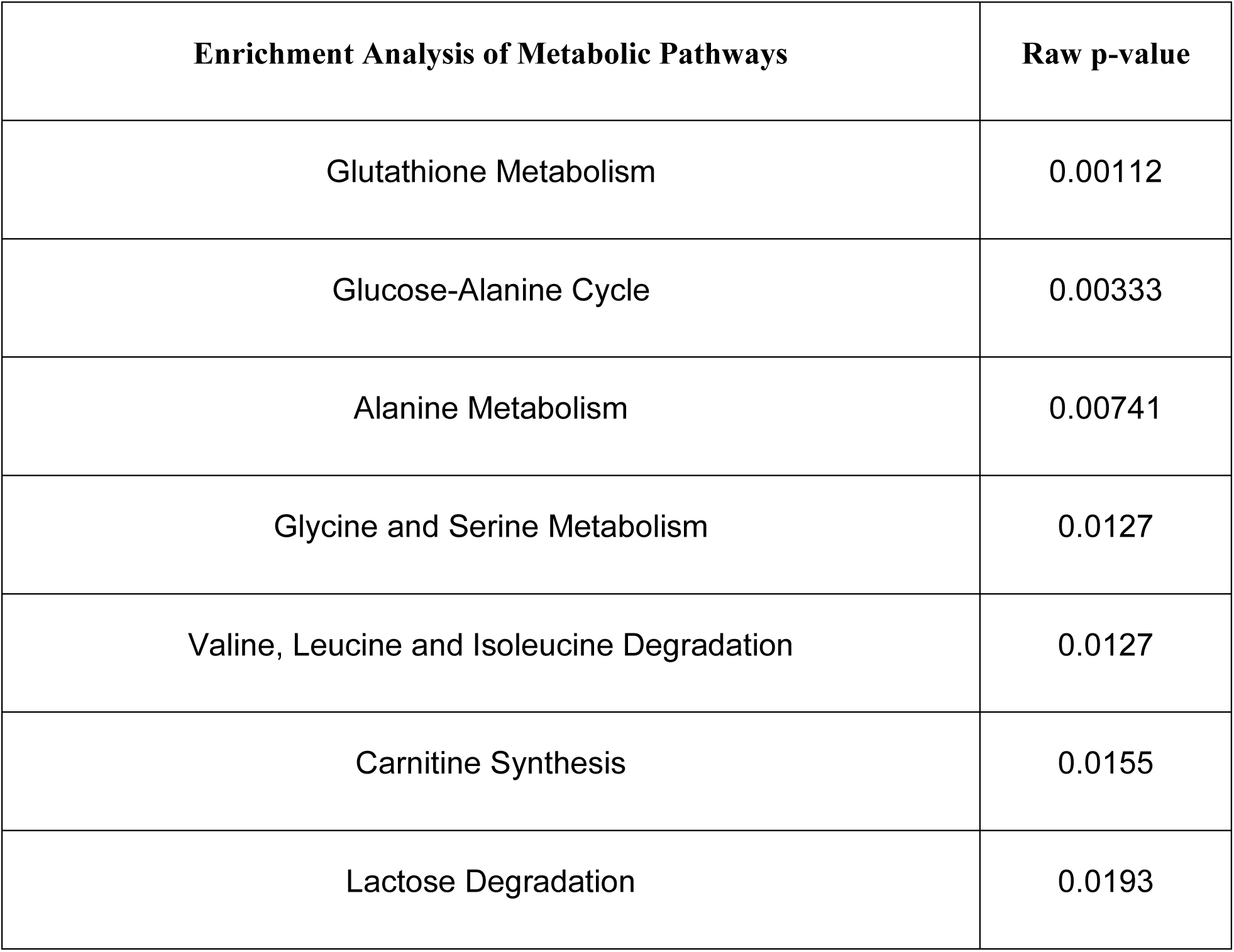
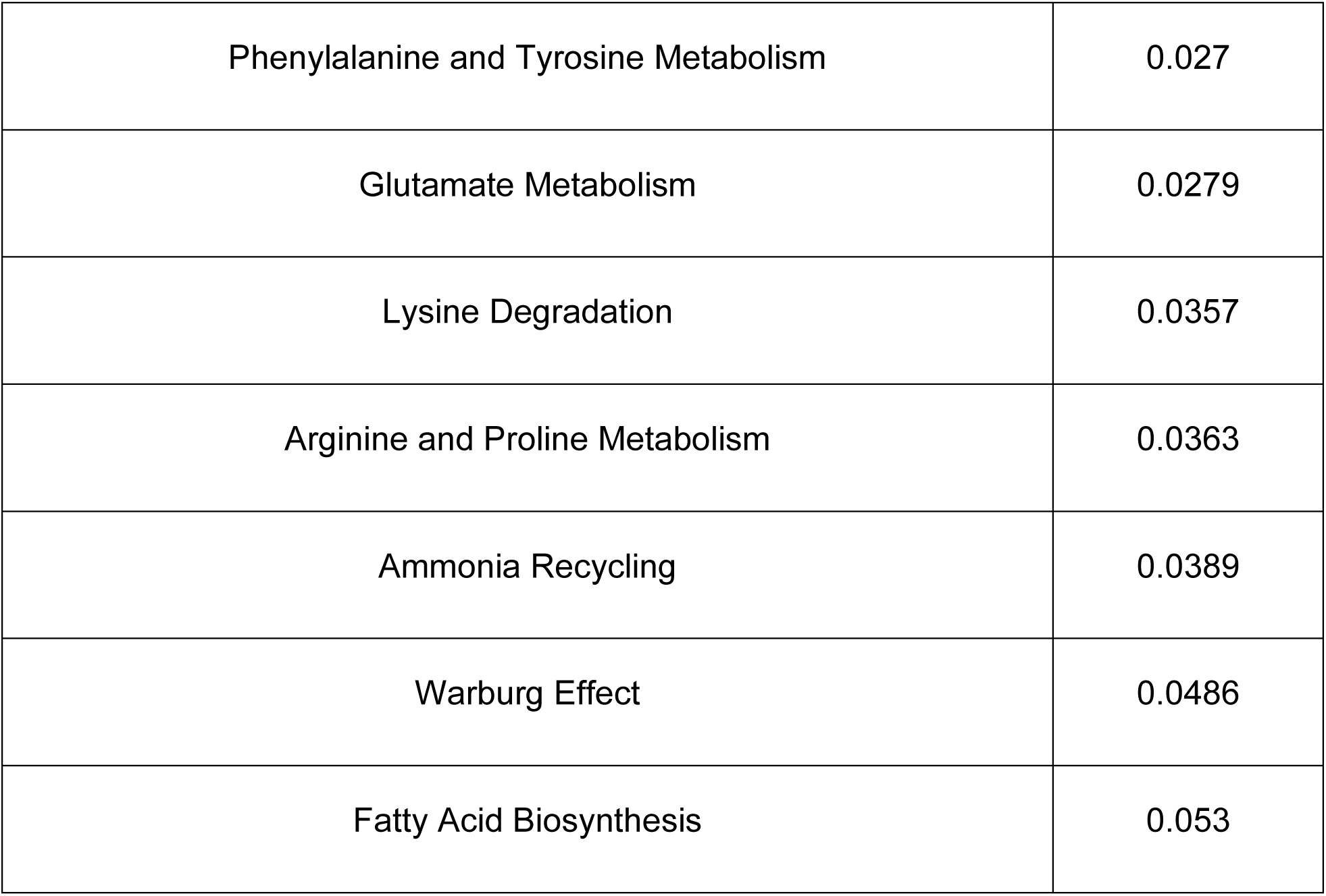
Enrichment analysis of metabolic pathways.

| Enrichment Analysis of Metabolic Pathways | Raw p-value |
| --- | --- |
| Glutathione Metabolism | 0.00112 |
| Glucose-Alanine Cycle | 0.00333 |
| Alanine Metabolism | 0.00741 |
| Glycine and Serine Metabolism | 0.0127 |
| Valine, Leucine and Isoleucine Degradation | 0.0127 |
| Carnitine Synthesis | 0.0155 |
| Lactose Degradation | 0.0193 |
| Phenylalanine and Tyrosine Metabolism | 0.027 |
| Glutamate Metabolism | 0.0279 |
| Lysine Degradation | 0.0357 |
| Arginine and Proline Metabolism | 0.0363 |
| Ammonia Recycling | 0.0389 |
| Warburg Effect | 0.0486 |
| Fatty Acid Biosynthesis | 0.053 |

**Table 2:** Impacted metabolic pathways in response to Trehalose.

| Metabolic Pathways Impacted | Raw p | Impact |
| --- | --- | --- |
| Valine, leucine and isoleucine biosynthesis | 5.12E-06 | 0 |
| Phenylalanine, tyrosine and tryptophan biosynthesis | 0.001752 | 1 |
| Phenylalanine metabolism | 0.007827 | 0.35714 |
| Galactose metabolism | 0.010749 | 0.39113 |
| Glutathione metabolism | 0.011897 | 0.11548 |
| Alanine, aspartate and glutamate metabolism | 0.011897 | 0.19712 |
| Glyoxylate and dicarboxylate metabolism | 0.017185 | 0.11 |
| Glycine, serine and threonine metabolism | 0.018683 | 0.4744 |
| Biosynthesis of unsaturated fatty acids | 0.023603 | 0 |
| Butanoate metabolism | 0.027199 | 0 |
| Valine, leucine and isoleucine degradation | 0.031165 | 0 |
| Neomycin, kanamycin and gentamicin biosynthesis | 0.034877 | 0 |
| Pantothenate and CoA biosynthesis | 0.04663 | 0 |
| Fatty acid biosynthesis | 0.047128 | 0.01473 |
| One carbon pool by folate | 0.074782 | 0.05637 |
| Lysine degradation | 0.095904 | 0.11247 |

## Discussion

### Trehalose reveals a mitochondria-linked vulnerability and selectively cooperates with mitochondrial-active therapies

Trehalose exhibits anti-inflammatory and anticancer properties and can be used in combination with standard anti-cancer drugs that have abilities to modulate mitochondria-associated parameters in tumour cells. This study identifies trehalose as a modulator of mitochondrial function in breast cancer and shows that its therapeutic interactions are not broadly cytotoxic but selectively synergistic with agents that intersect mitochondrial physiology and cellular structural stress (tamoxifen, colchicine; moderate with 2-DG; subtype-dependent with paclitaxel). The predominance of synergy in MCF7 relative to MDA-MB-231 suggests that trehalose exploits a vulnerability enriched in ER⁺ cells—namely, higher reliance on mitochondrial bioenergetic capacity and mitochondrial maintenance programs.

### Trehalose imposes mitochondrial dysfunction, checkpoint activation, and a pseudo-hypoxic stress program in ER⁺ cells

Trehalose is known for its autophagy-inducing actions and its anti-inflammatory actions. Similar to these effects, trehalose reduced ROS and MMP and produced a cytochrome c–responsive increase in respiration, indicating compromised mitochondrial membrane integrity. Together with S/G2-M accumulation, CDK4 suppression, and p21 induction, these results support a model where trehalose triggers bioenergetic stress sufficient to activate growth-restrictive checkpoints. The accompanying increase in HIF-1α with suppression of VEGFA is consistent with a pseudo-hypoxic programme that signals metabolic restraint without initiating a canonical pro-angiogenic switch. Increased EMT/stemness-associated markers likely reflect stress-adaptive transcriptional remodeling rather than a straightforward metastasis activation; importantly, this phenotype appears coupled to growth restriction and mitochondrial suppression in this system [39].

### Trehalose suppresses mitochondrial maintenance/biogenesis and enforces mitochondria-dependent cell death

Trehalose reduced mitochondrial abundance and suppressed TFAM, supporting impaired mtDNA maintenance and mitochondrial renewal. The enhancement of caspase processing by trehalose, especially in combination with tamoxifen and colchicine, indicates reinforcement of mitochondria-linked apoptosis. The decisive observation that trehalose fails to reduce viability or respiration in MCF7 ρ⁰ cells establishes mitochondrial dependency as a core mechanistic feature. Thus, trehalose is best understood as a mitochondrial state modulator that converts mitochondrial reliance into a liability in ER⁺ cells.

### Subtype-specific bioenergetic architecture explains differential trehalose sensitivity

Functional respirometry showed that trehalose constrains ETS capacity and impairs complex I–linked respiration in MCF7 cells, increasing L/E indices due to reduced maximal electron transport. In MDA-MB-231 cells, high baseline uncoupling and preserved membrane integrity likely buffer the impact of trehalose on mitochondrial flux. These findings align with transcriptomic patterns showing stronger respiratory programmes in ER⁺ contexts and with clinical survival associations linking elevated respiratory gene expression to poorer outcomes in breast cancer cohorts. The metabolic dependency on mitochondria in MCF7 cells is no longer valid in MDA-MB-231 cells. These findings correlate clearly define well established notion of metabolic heterogeneity in BC. Collectively, these data support a framework in which OxPhos dependency and mitochondrial programme maintenance determine trehalose responsiveness.

### A BCAA–glycolysis–lipid axis defines the dominant metabolic reconfiguration downstream of mitochondrial constraint

In proliferating cancer cells, amino acids are metabolically linked to glycolysis via transamination, supporting TCA replenishment, biosynthesis, and redox balance; accordingly, amino acid restriction suppresses tumour growth [40]. Importantly, the findings indicate a coordinated reduction rather than selective BCAA accumulation or diversion, arguing against compensatory amino acid oxidation and supporting global suppression of growth-linked carbon–nitrogen integration [41, 42, 43]. Beyond reduced BCAA flux, serine and glutamate levels were elevated in trehalose-treated ER/PR⁺ breast cancer cells, in contrast to prior TNBC studies where these amino acids are preferentially increased, suggesting metabolic differences [44]. The increased serine proportion may reflect hyperactivation of serine biosynthesis, as trehalose reduces lactate production and likely diverts early glycolytic flux toward serine anabolism. In contrast, elevated glutamate is well established to be associated with CD8⁺ T-cell activation [45, 46].

De novo lipogenesis and lipid metabolism have contrast meaning in terms of different BC subtypes. In TNBC, lipid metabolism and de novo biosynthesis are hyperactivated, while ER/PR+ve BC cells have more controlled de novo biosynthesis [47, 48]. The exceptionally strong positive correlation between oleic acid and cholesterol, together with their negative association with BCAAs, suggests that lipid accumulation represents an actively maintained metabolic state rather than a passive consequence of reduced proliferation. This is further supported by enrichment of fatty acid biosynthesis and sphingolipid-related pathways [49]. Lipid accumulation might emerge as an alternative metabolic state rather than a secondary consequence, including membrane structure and fluidity as well as lipotoxicity [50, 51, 52].

The current study supports this scenario, but also specifies a different set of metabolic patterns where, along with reduced mitochondrial biogenesis and OxPhos, high lipid storage and biosynthesis correlate with a significant decrease in amino acid biosynthetic pathways. This particular metabolic pattern is a stable lipid-centric metabolic reprogramming associated metabolic rewiring, followed by differential activation of catabolic or anabolic pathways, which may also be the case with trehalose-associated metabolic stress, and thus induce metabolic rewiring [53].

Metabolomics demonstrated that trehalose does not induce a non-specific collapse but rather establishes a coordinated metabolic state characterized by reduced glycolytic output (lower lactate), depletion of BCAAs/amino acids, and accumulation of long-chain fatty acids and cholesterol (**Fig. 5E**). Multivariate feature selection and correlation structure converge on a dominant metabolic axis: a coupled BCAA–glycolysis module that is inversely related to a lipid-accumulation module. This organization is consistent with reduced anabolic/growth signalling output (including attenuation of PI3K/Akt) and insufficient compensatory restoration of oxidative capacity, resulting in a lipid-buffered, bioenergetically constrained state.

### Conclusion

Across pharmacologic interactions, mitochondrial physiology, cell-cycle control, mitochondrial biogenesis markers, ρ⁰ dependency, transcriptomic context, and metabolomic state mapping, trehalose consistently enforces a mitochondrial-constrained phenotype in ER⁺ breast cancer. The most parsimonious model is that trehalose limits mitochondrial maintenance and maximal ETS capacity, suppresses growth-linked carbon–nitrogen integration (BCAA biosynthesis), and stabilizes a lipid-enriched metabolic state, thereby sensitizing ER⁺ cells to mitochondrial-active therapies such as tamoxifen and colchicine. Across all analytical layers, trehalose exposure produced a consistently different metabolic pattern displaying the aggressive metabolic plasticity of ER+ve BC cells, a finding that warrants further research involving both preclinical and clinical models.

### Limitations of the Study

These findings were generated in vitro and do not capture tumour microenvironment constraints, immune interactions, or systemic metabolic effects. Additional work is needed to define the proximal molecular target(s) of trehalose within mitochondrial biology, to establish in vivo relevance (including pharmacokinetics and achievable tissue exposures), and to determine whether the trehalose-induced pseudo-hypoxic/stress programme yields durable phenotypic consequences (including EMT/stemness markers) in tumour models. The downstream role of c-Jun and broader transcriptional networks also warrant deeper mechanistic evaluation.

## Materials and Methods

### Cell Culture

The different cell lines for breast cancer [MCF-7 and T47D (ER+), MDA-MB-231 (TNBC)], lung cancer (A549), neuroblastoma (SH-SY5Y) were purchased from the National Cell Repository, India. The human PBMCs (hPBMCs) were isolated from healthy individual for using them as non-cancerous cells. Cells were cultured in DMEM supplemented with 10% FBS (fetal bovine serum) and Penicillin/Streptomycin cocktail (1X) and incubated in the CO2 incubator with 5% CO2 at 37**°**C. Cells were seeded into 96-well plate for analyzing the cytotoxicity properties of investigational drugs.

### Combenefit and computer-based combinatorial studies

The cancer cell lines MCF7, MDA-MB-231, T47D, SHSY5Y, and A549 were purchased from NCCS, Pune. The cells were grown in complete DMEM media, and an MTT assay was performed to analyse the cytotoxicity of different drugs. [54]. In combinatorial therapy studies, the concentrations of Trehalose used were 0.1mM, 0.5mM, 1mM, and 5mM, while 1 µM, 2.5 µM, 5 µM, and 10 µM were used for other standard drugs. For the 2-DG, the concentration used were 1mM, 2.5mM, 5mM and 10mM. The Trehalose and all other standard drugs in combination were given for 48 hours. The % cell cytotoxicity values were used in analysing the quantitation of synergism and antagonism between two drugs using software, i.e., compusyn and Combenefit (**Supplementary Data**).

### Flow Cytometry-based analysis of cellular parameters

The MCF7 cells were trypsinised, washed with PBS, and incubated with H2DCFDA dye at 10µM final concentration and JC-1 dye at 5µM final concentration for 60 min in the dark at RT. After washing the cells with PBS, the cells were analysed for emission spectra using a BD-Accuri C6 plus flow cytometer. [54]. The MCF7 cells were trypsinised and washed with PBS. For cell cycle analysis, serum starvation before treatment and post-treatment fixation with 70% ethanol for 3-4 hours at 4°C were performed. The cells were incubated with respective dyes for the cell cycle (PI/RNAse solution) and PI/Annexin-V assay (PI solution and Annexin-V FITC bound) as per the manufacturer’s protocol. The cells were analysed using a BD Accuri C6 flow cytometer.

### Quantitative real-time PCR

The MCF7 cells were trypsinised, and RNA isolation was performed using the Trizol method. The cDNA was synthesised using the Revert-Aid cDNA synthesis kit. The cDNA was diluted 5 times, and qPCR was performed as per the manufacturer’s protocol (SYBR & CFX-96 qPCR machine, Bio-Rad). The primers used in the study are listed in supplementary files (**Fig. S11**).

### Immunoblotting

The MCF7 cells were trypsinised and lysed using RIPA lysis. Protein estimation was done by Bradford assay, and proteins were separated by electrophoresis on 8–15% SDS-PAGE and blotted onto a nitrocellulose membrane. Intermediate washing of the membrane is done with TBST. Blocking was done in a 3% non-fat milk solution and incubated with the primary antibody. Further, incubated for 1 h with an HRP-conjugated secondary antibody.

The immune detection was carried out and visualised with chemiluminescence (Bio-Rad, USA). The data was normalised to the proteomic expression level of β-Actin.

### Generation of Rho-0 Cells and Live/Dead Cell Assay

The MCF7 cells were treated with ethidium bromide (50 ng/ml) for 6–8 weeks for the depletion of mitochondrial DNA (mtDNA) [55]. The MCF7-Rho-0 cells (ρ0 cells) were cultured in complete DMEM supplemented with 50 mg/ml uridine. [55]. The cells were treated with Trehalose in DMEM media without uridine supplementation and were trypsinised. The appropriate concentration of calcein AM and ethidium homodimer-1 in PBS was added as per the manufacturer’s protocol (Cytotoxicity Kit-Molecular Probes^TM^) and incubated for 30 minutes at RT. The live and dead cells were analysed using an Attune Nxt flow cytometer (Thermofisher Scientific).

### Assessment of Mitochondrial Mass

The cells cultured on coverslip were stained with Mitotracker Green^TM^ (250nM), Invitrogen, and kept in the dark for 45 minutes at room temperature. Further, the cells were fixed with acetone: methanol (1:1) for 10 minutes and then washed with PBS. The cells were rewashed with PBS, and a coverslip was placed on the slide using fluoro-shield mounting media and kept in the dark. The slides were analysed using a confocal microscope (Olympus FV-1200).

The cells cultured without a coverslip were trypsinised and incubated in the dark with 250 nM Mitotracker Green^TM^ for 45 minutes at room temperature. The cells were centrifuged and resuspended in PBS for flow cytometric analysis (BD Accuri C6 flow cytometer) [56].

### GEO Database Analysis

The GEO database with accession IDs including GSE215917, GSE174152, GSE68815, GSE219274, GSE223350, GSE234171, and GSE188914 was selected based on therapies used and data availability. The expression of key mitochondria encoded genes including ND1, ND2, ND3, ND4, ND4L, ND5, ND6, SDHA, SDHB, SDHD, CYB, CYC1, COX1, COX2 and COX3, ATP6 and ATP8 was analyzed.

### High-Resolution Respirometry-based SUIT protocols for assessing mitochondrial parameters

#### Basal OCR analysis

The MCF7 and MDA-MB-231 cells were trypsinised and resuspended in 2 mL DMEM media with a cell density of 10^6^ cells/mL. The basal OCR consumption and O2 concentration were measured for 20 minutes until the oxygen flux became constant. The calibrated slope readings for O2-flux and O2 concentration from respective DL-protocol were taken to plot the graph (O2k Fluo-respirometer, Austria) [56, 57, 58].

#### Mitochondrial membrane integrity analysis

For mitochondrial membrane integrity analysis, digitonin (12µM) was added to permeabilise the cells (state-2), and oxygen consumption is linked to residual oxygen consumption rate. The succinate (5mM) was added before the ADP substrate (1mM) was added to achieve state-3. The Cytochrome C (0.5µM) was added to analyse the oxygen consumption rate (OCR) [57, 58].

#### Leak respiration, ETS capacity and Coupling efficiency analysis

For coupling/uncoupling efficiency analysis, oligomycin (5nM) was added to analyse leak respiration (L-state). The FCCP titration (2µM) was done to achieve the respiratory unit’s ETS capacity (E-state). The rotenone (0.5µM), i.e., complex I inhibitor, and Antimycin A (2.5µM), i.e., complex III inhibitor, were added to analyse the residual oxygen consumption rate. The LEAK/ETS respiration ratio was calculated to analyse coupling/uncoupling respiration.

#### Activity of mitochondrial respiratory complexes

For analyzing the activity of mitochondrial respiratory complexes, digitonin (12µM) was added to permeabilise the cells, then succinate (10mM) was added without adenylate contributes to S-linked LEAK respiration; ADP (2.5mM) addition refers to Routine respiration: Pyruvate (5mM) addition induce complex I activity and linked to Routine respiration; Rotenone (0.5µM) inhibit complex I and O2 consumption is linked to complex II-mediated respiration. Antimycin A inhibits complex III, and O2 consumption is linked to non-mitochondrial respiration.

#### Metabolite Extraction and GC–MS/MS Analysis

MCF7 breast cancer cells were treated with low- and high-dose trehalose for 48 hours in triplicate, and cells were washed with sterile phosphate-buffered saline (PBS). Metabolism was quenched with 1 mL of ice-cold methanol:water (80:20) per plate and incubated on ice for 10 min. Cells were scraped, transferred to sterile tubes, and centrifuged at 2,500 rpm at 4 °C for 10 min. The metabolite-containing supernatant was collected and stored at −80 °C.

For GC–MS/MS analysis, 200 µL of each extract was transferred into glass vials, and 20 µL of L-norvaline was added as an internal standard. Samples were dried under nitrogen gas and derivatised using N-methyl-N-(trimethylsilyl) trifluoroacetamide (MSTFA) with dimethyl sulfoxide (DMSO), followed by incubation at 80 °C for 30 min. Derivatised samples were analysed on a GC-MS-TQ8050 NX system (Shimadzu, Japan) operated in electron-impact ionisation mode. Mass spectra were acquired over the 50–600 m/z range, and metabolites were identified using GC-MS Solution software with reference to the NIST library [59, 60].

#### Metabolite Curation and Pathway Analysis

All detected metabolites were manually curated based on biological plausibility, retention index consistency, derivatisation chemistry, and comparison against DMEM background controls to exclude exogenous contaminants. High-confidence metabolites were mapped to curated human metabolic pathways using KEGG and HMDB databases. Pathway enrichment was performed using a hypergeometric distribution model, with P < 0.05 considered statistically significant. Hierarchical clustering and heatmap visualisation were conducted using z-score–normalized metabolite intensities [59, 60].

#### Statistical Analysis

The graphs were plotted, and the statistics were applied using GraphPad Prism (v8.0). The results were considered significant with p<0.05 and are provided as means ± SEM for experiments done in triplicate. The group values were expressed as a two-way analysis of variance (ANOVA). They were followed by Dunnett’s post hoc multiple comparison test and Sidak’s post hoc multiple comparison test as per the data input using GraphPad Prism v8.0. Metabolite peak areas were averaged from biological triplicates, normalised, log-transformed, and autoscaled. PCA and PLS-DA were performed using MetaboAnalyst 6.0, and Variable Importance in Projection (VIP) scores were calculated. Only metabolites meeting a high VIP score and Pearson correlation value (r ≥ 0.95) were considered high-confidence metabolic regulators. Pearson correlation analyses were conducted in MetaboAnalyst and independently validated using GraphPad Prism v8.0 [59, 60].

## Declaration

### Author’s Contribution

TS and PK performed the experiments, TS & PK analysed the data, compiled results, and wrote the manuscript. AM and SS conceived the study and participated in data analysis and writing. All the authors have reviewed and approved the manuscript.

## Acknowledgement and funding

TS is thankful to CSIR/UGC for financial assistance (grant no. 09/1051(0042)/2019-EMR-I). The authors thank the DST-FIST grant (SR/FST/LS-I/2017/49) to the Department of Human Genetics and Molecular Medicine, Central University of Punjab. The authors acknowledge the Central University of Punjab for providing infrastructure support and funds for Oroboros O2K. SS and TS thank the DST-SERB EMR grant (SR/SO/AS-31/2014) for financial support. All authors are thankful to the equipment infrastructure provided by the Central Instrumentation Laboratory (CIL), Central University of Punjab, Bathinda, and the technical assistance by Dr Sumeer Razdan, Mr. Vikas, and Mr. Ajit Paul Singh of CIL, CUPB.

## Competing Interests

The authors declare no conflict of interest.

## Availability of data and material

The data analysed are provided in supplementary data, and raw data can be obtained upon request via email to TS.

## Research involving human participants and animals

This work includes no tissue/blood samples of humans or animals.

## Ethics approval and Consent to participate

Not applicable

## Consent for publication

All authors agreed to publish this article.

## Abbreviations

BC: Breast Cancer
ER/PR^+ve^: Estrogen/Progesterone-positive
TNBC: Triple Negative Breast Cancer
OxPhos: Oxidative Phosphorylation
ETC: Electron Transport Chain
Treh: Trehalose
Treh^Low^: 100µM Trehalose treated cells (Low Dose)
Treh^High^: 5mM Trehalose treated cells (High Dose)
MCF7^Treh^: Trehalose-treated MCF7 cells
CPT: Camptothecin
PTX: Paclitaxel
Dox: Doxorubicin
2-DG: 2-deoxyglucose
Treh-TF: Trehalose-Tamoxifen Combination
Treh-Col: Trehalose-Colchicine Combination
Treh-PTX: Trehalose-Paclitaxel Combination
Treh-2DG: Trehalose-2-dexoyglucose Combination
Treh-Er: Trehalose-Erlotinib Combination
Treh-Et: Trehalose-Etoposide Combination
Treh-CPT: Trehalose-Camptothecin Combination
Treh-Dox: Trehalose-Doxorubicin Combination
MCF7: In vitro ER/PR^+ve^ BC model
MDA-MB-231: In vitro TNBC model
TFAM: Mitochondrial transcription factor A
PGC-1α: Peroxisome proliferator-activated receptor gamma coactivator 1-alpha
CDK: Cyclin-Dependent Kinases
AMPK: AMP-activated protein kinase
L/E Ratio: Leak Respiration/ETC Ratio
L/P Ratio: Leak/Routine Respiration
ETC: Electron Transport Chain
BCAA: Branched-Chain Amino Acids

## Notes

### Competing Interest Statement

The authors have declared no competing interest.

