## Supplementary Data for "Disruption of PI3K-OxPhos Coupling by Trehalose Drives a BCAA-to-Lipid Metabolic Switch in Hormone-Receptor–Positive Breast Cancer"

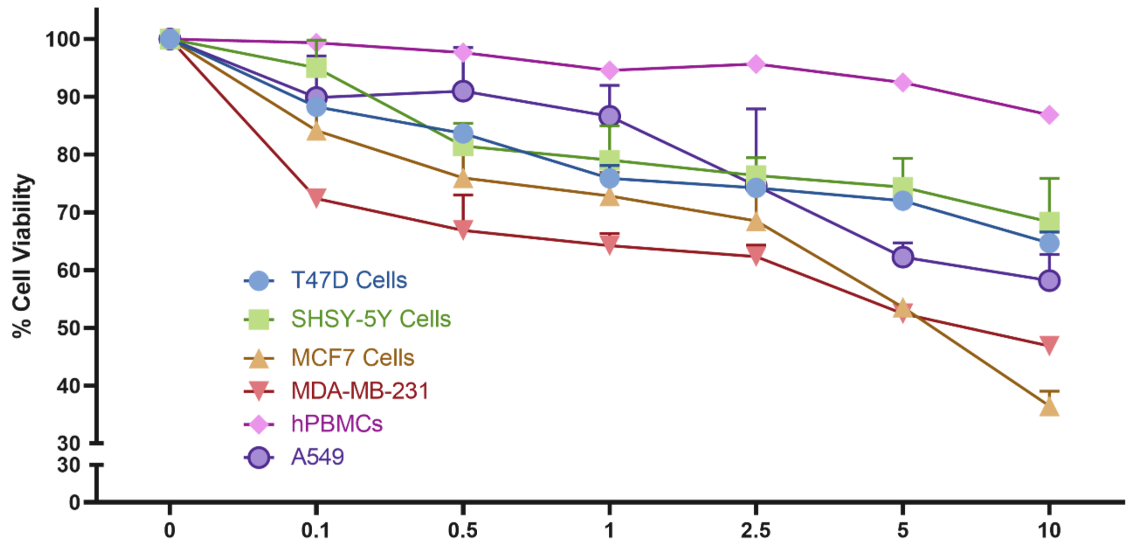

**Figure S1:** Dose-dependent antiproliferative potential of the Trehalose across cancer and non-cancer cell types. Cell viability (%) was assessed in T47D ($ER^{+ve}$ BC), MCF7 ($ER^{+ve}$ BC), MDA-MB-231 (TNBC), SH-SY5Y (neuroblastoma), A549 (lung adenocarcinoma), and human peripheral blood mononuclear cells (hPBMCs) following treatment with increasing concentrations of the trehalose (0–10 µM). Cell viability is expressed as percentage relative to untreated control (0 µM), which was set to 100%. Data are presented as mean ± SD from independent experiments.

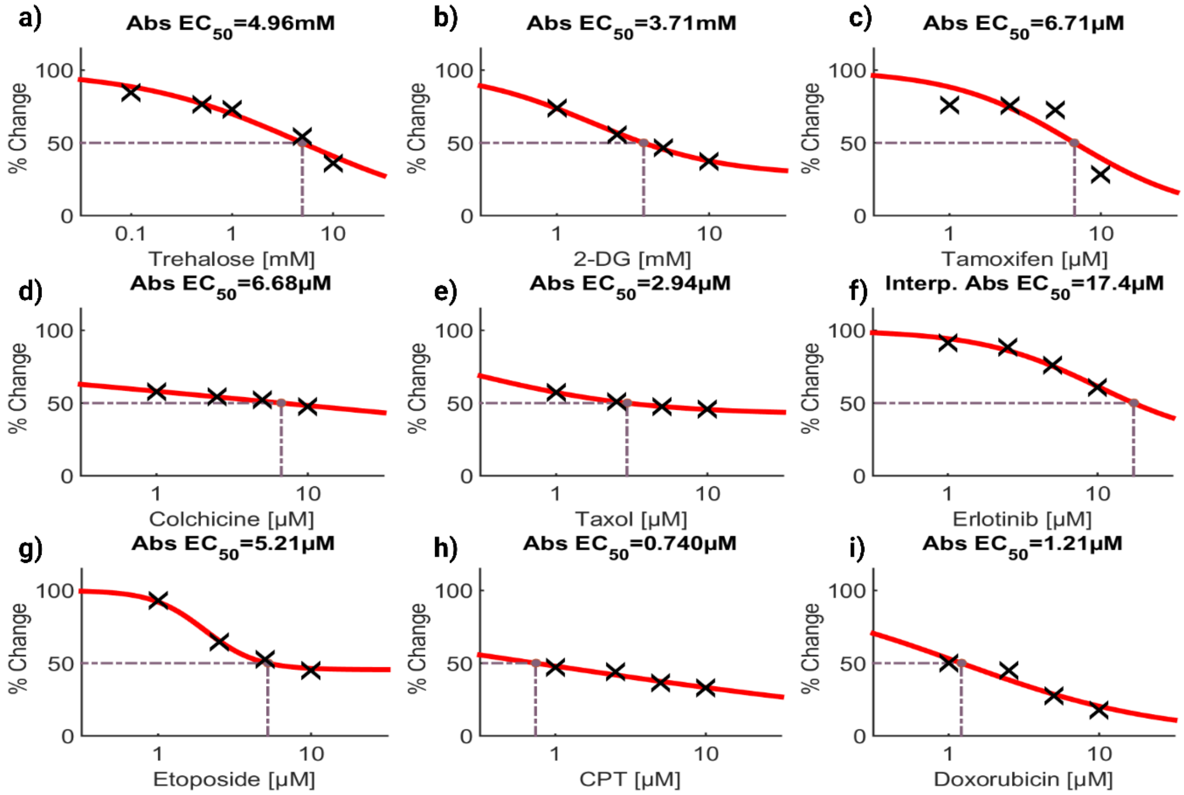

**Figure S2:** The $IC_{50}$ of Trehalose, Tamoxifen, Colchicine, Paclitaxel (Taxol), 2-deoxy glucose, camptothecin, erlotinib, etoposide and doxorubicin in a) MCF7 cells calculated using HSA method in combenefit software.

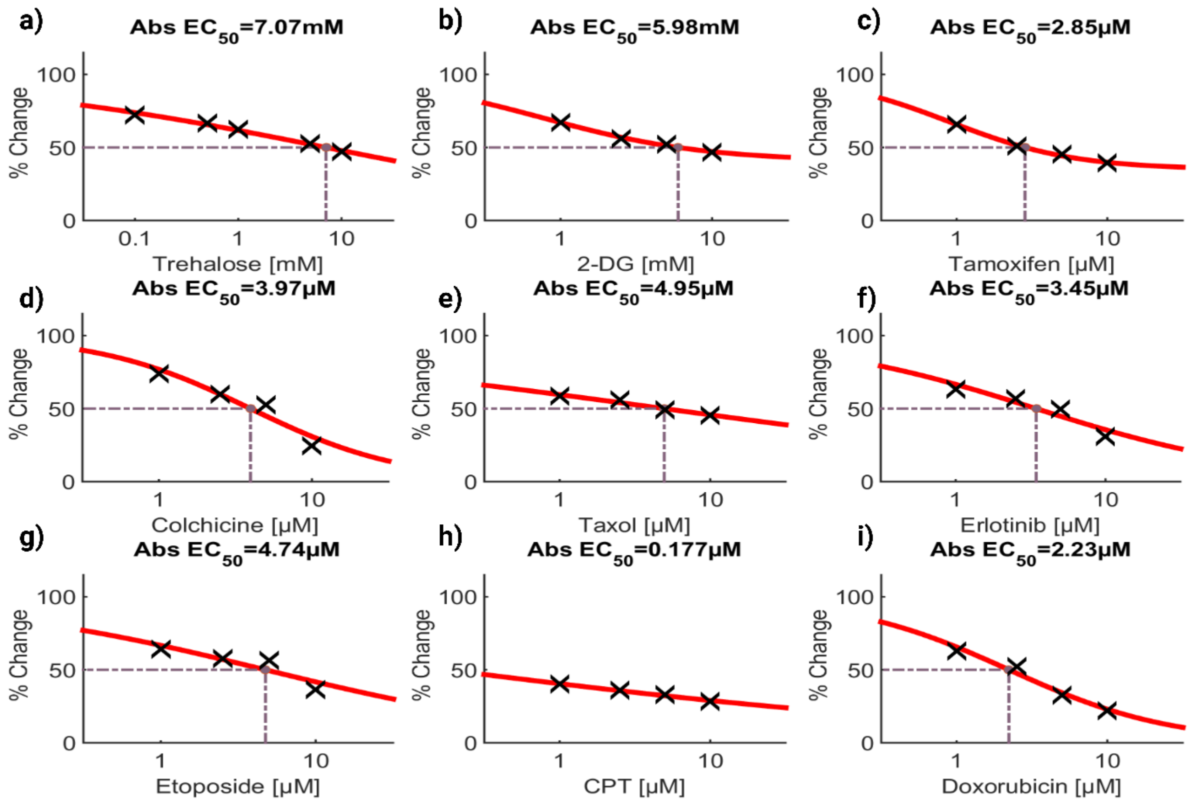

**Figure S3:** The $IC_{50}$ of Trehalose, Tamoxifen, Colchicine, Paclitaxel (Taxol), 2-deoxy glucose, camptothecin, erlotinib, etoposide and doxorubicin in MDA-MB-231 cells calculated using HSA method in combenefit software.

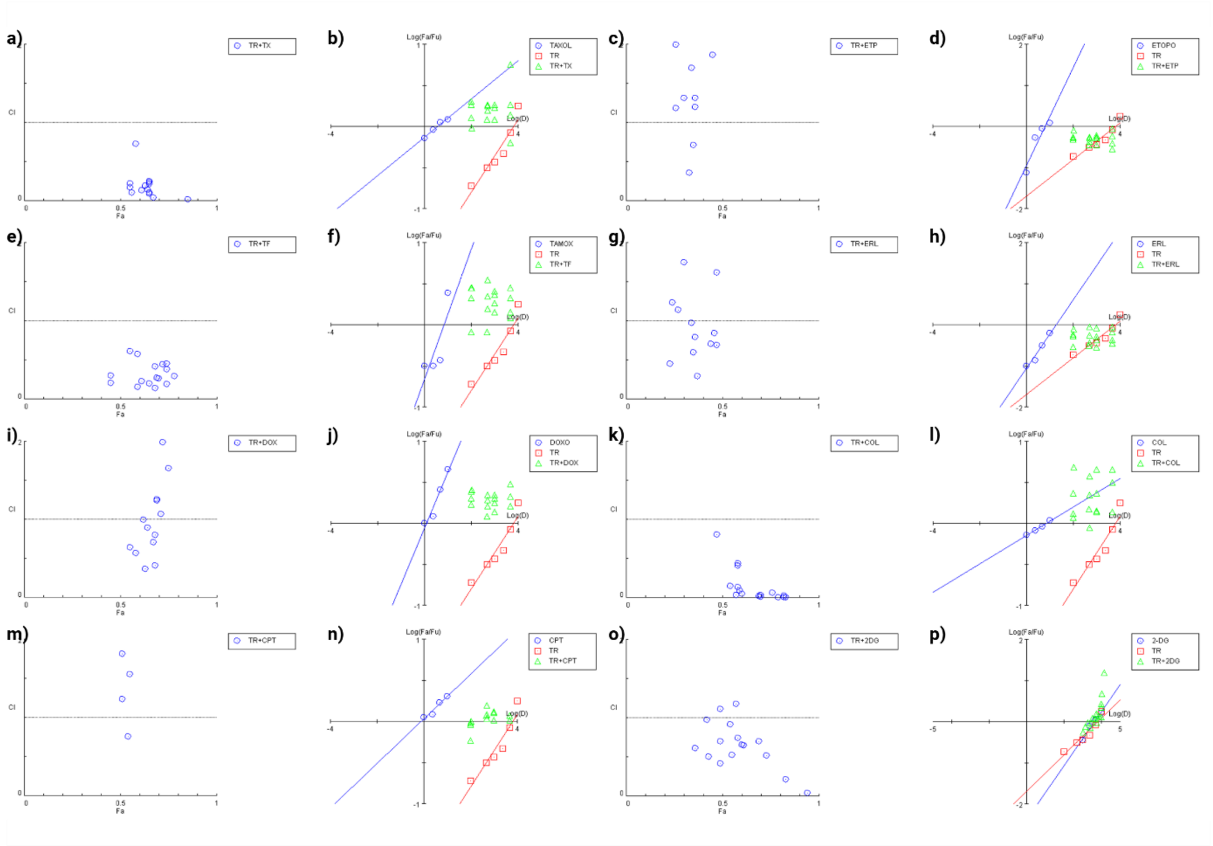

**Figure S4:** The synergistic or antagonistic cytotoxic impact of trehalose in combination with different drugs on MCF7 cells using HSA (Highest Single Agent) based compusyn software analysis. The combination index plotted CI values close to zero (near the x-axis) and less than 1, which signifies the combination to be synergistic, while more than 2 signifies the antagonistic effects for given combinations. The combination index plot for trehalose in combination with a) Taxol, c) etoposide, e) tamoxifen, g) erlotinib, i) doxorubicin, k) colchicine, m) etoposide, and o) 2-deoxyglucose (2-DG). The median effect plot for trehalose in combination with b) Taxol, d) etoposide, f) tamoxifen, h) erlotinib, j) doxorubicin, l) colchicine, n) etoposide, and p) 2-deoxyglucose (2-DG).

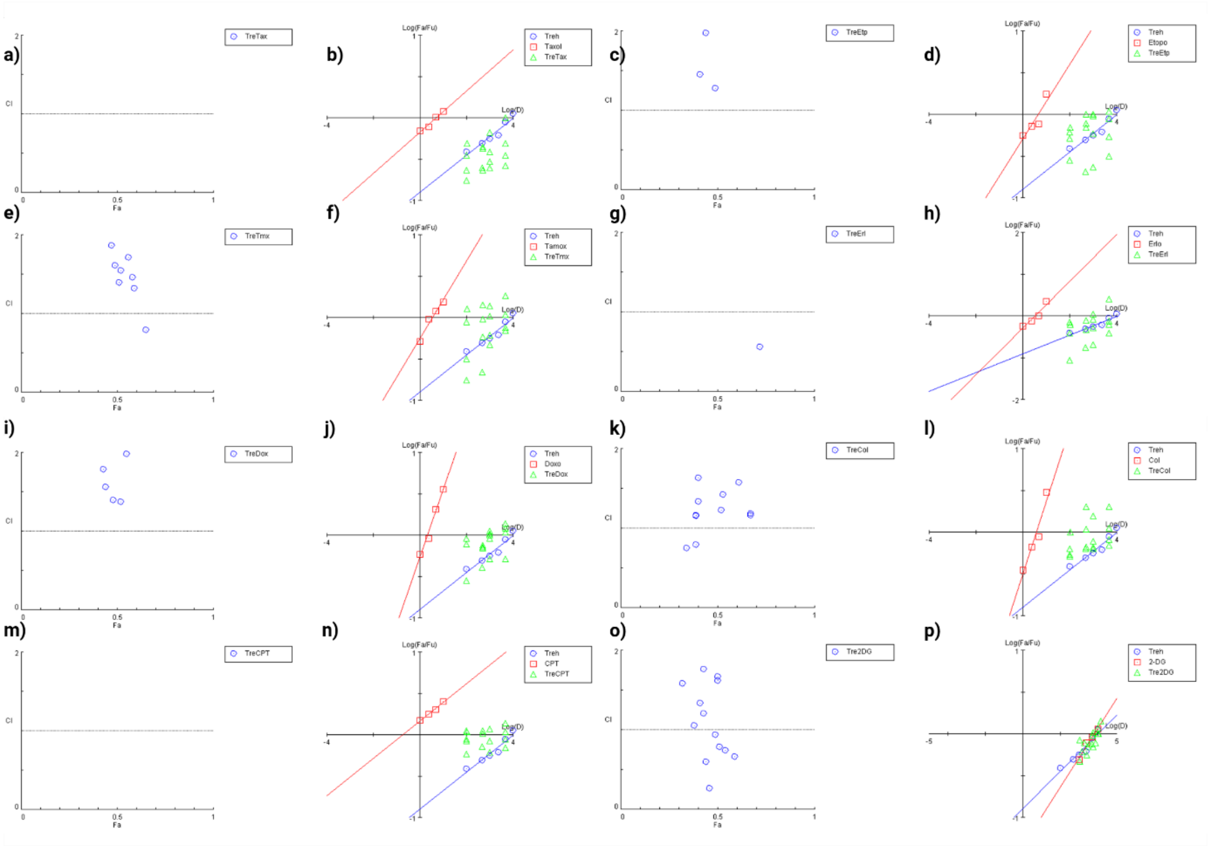

**Figure S5:** The synergistic or antagonistic cytotoxic impact of trehalose in combination with different drugs on MDA-MB-231 cells using HSA (Highest Single Agent) based compusyn software analysis. The combination index plot for trehalose in combination with a) Taxol, c) etoposide, e) tamoxifen, g) erlotinib, i) doxorubicin, k) colchicine, m) etoposide, and o) 2-deoxyglucose (2-DG). The median effect plot for trehalose in combination with b) Taxol, d) etoposide, f) tamoxifen, h) erlotinib, j) doxorubicin, l) colchicine, n) etoposide, and p) 2-deoxyglucose (2-DG).

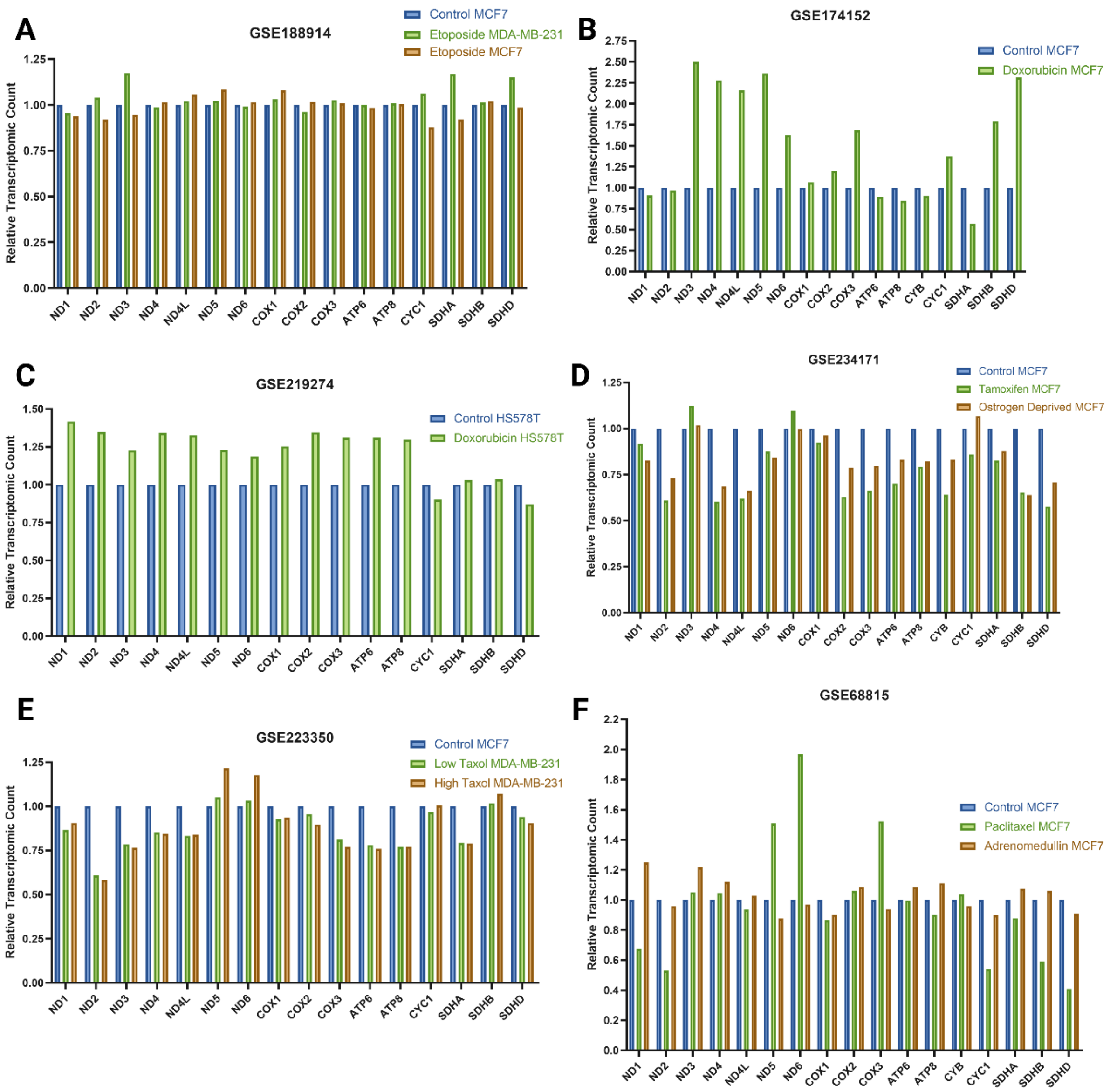

**Figure S6:** The GEO2R based expression of mitochondrial encoded genes in breast cancer subtypes under normal and therapeutic conditions.

**(A)** Expression analysis of mitochondrial encoded metabolic genes in MCF7 and MDA-MB-231 cells treated with etoposide using RNA-seq data.

**(B)** Expression analysis of mitochondrial encoded metabolic genes in MCF7 cells treated with doxorubicin (Dox) using RNA-seq data.

**(C)** Expression analysis of mitochondrial encoded metabolic genes in HS578T cells treated with doxorubicin (Dox) using RNA-seq data.

**(D)** Expression analysis of mitochondrial encoded metabolic genes in MCF7 cells treated with tamoxifen (TF) and oestrogen deprivation (OD) using RNA-seq data.

**(E)** Expression analysis of mitochondrial encoded metabolic genes in MDA-MB-231 cells treated with low and high doses of paclitaxel (PTX) using RNA-seq data.

**(F)** Expression analysis of mitochondrial encoded metabolic genes in MCF7 cells treated with paclitaxel (PTX) and adrenomedullin (ADM) using RNA-seq data.

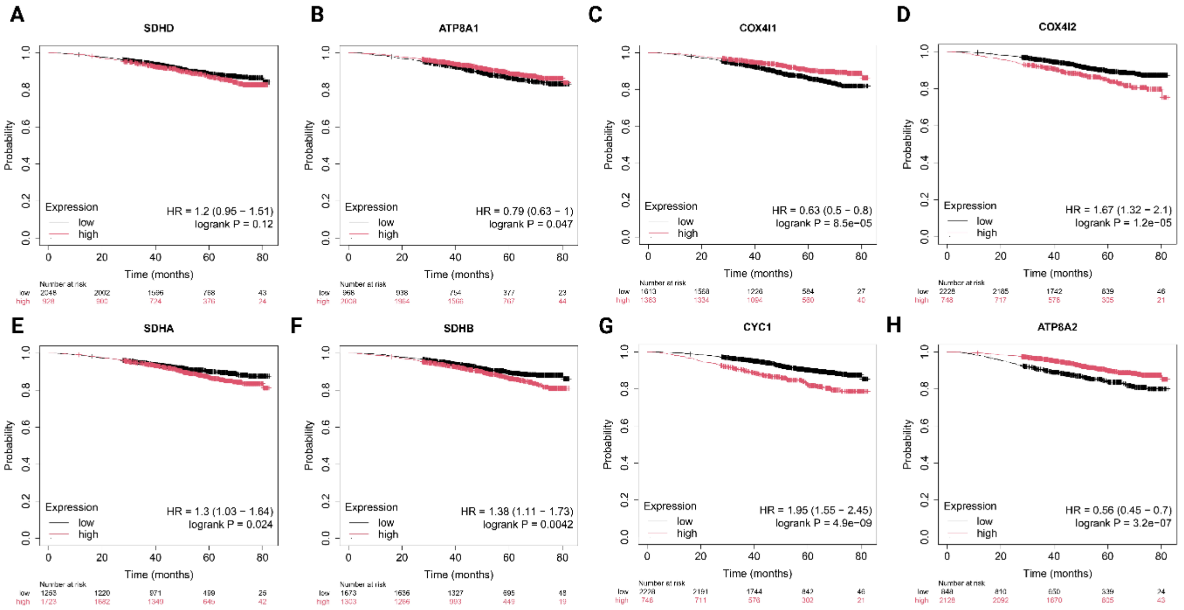

**Figure S7:** Kaplan-Meirr Plot for key mitochondrial encoded genes in association with progression of breast cancer and survival of patients among BC patients.

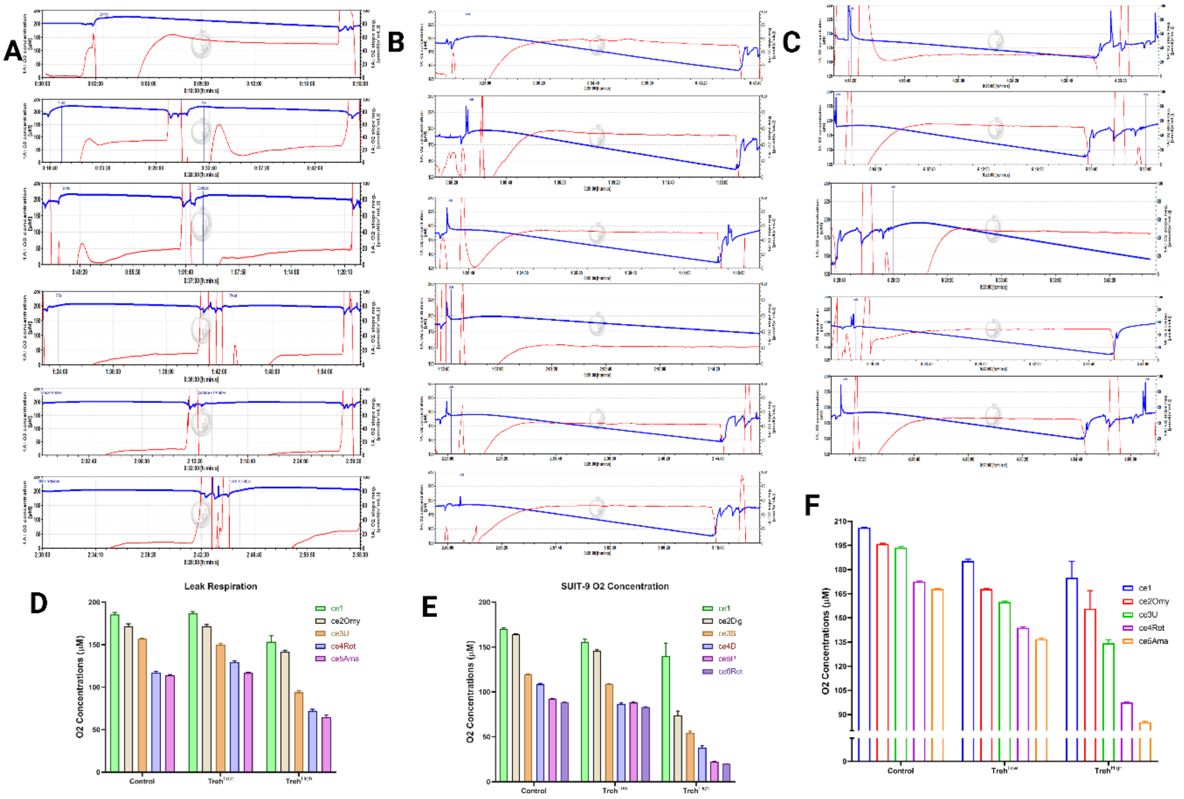

**Figure S8:** Combinatorial drug response-based OCR of Trehalose, effective chemotherapeutics and the combination of Trehalose with these screened drugs in MCF7 and MDA-MB-231 cells.

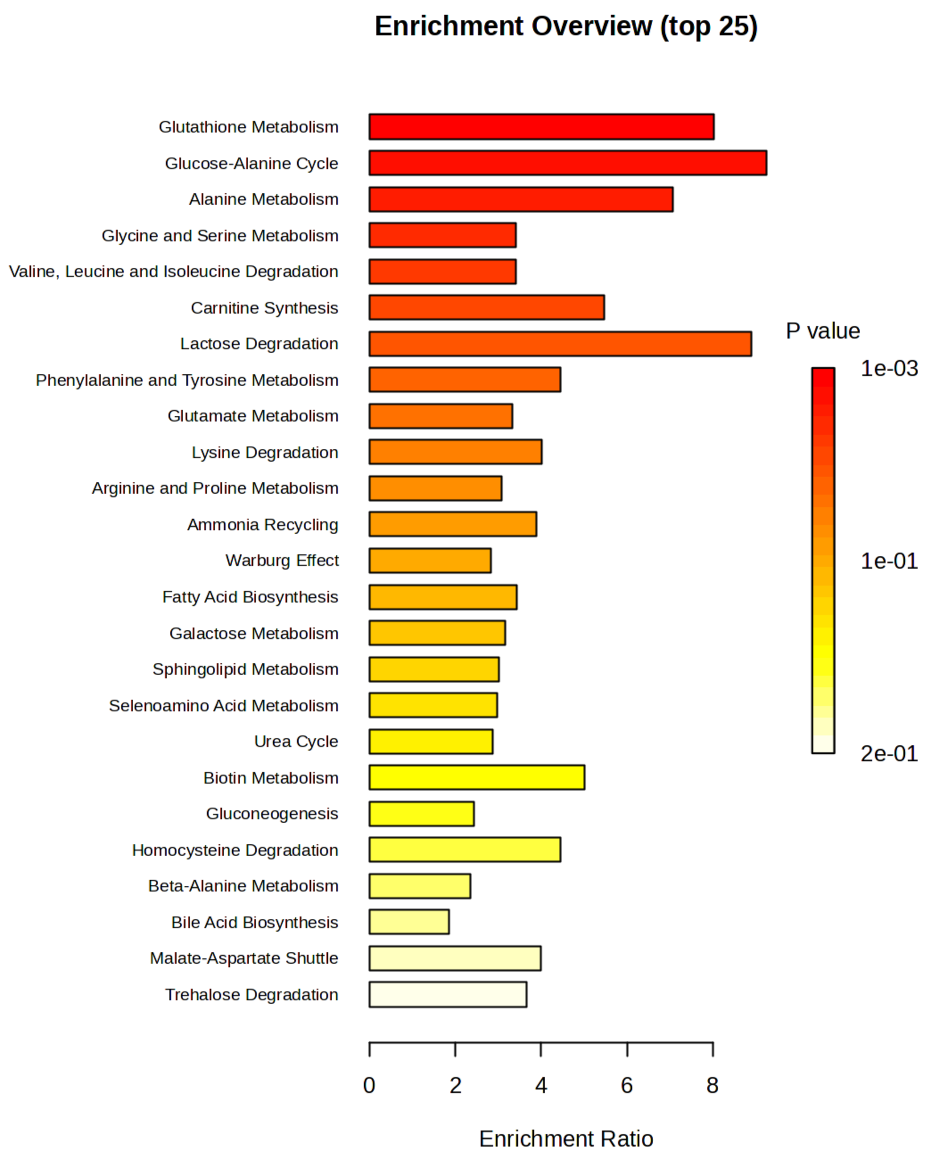

**Figure S9:** Pathway enrichment analysis of key metabolites altered under Trehalose treatment.

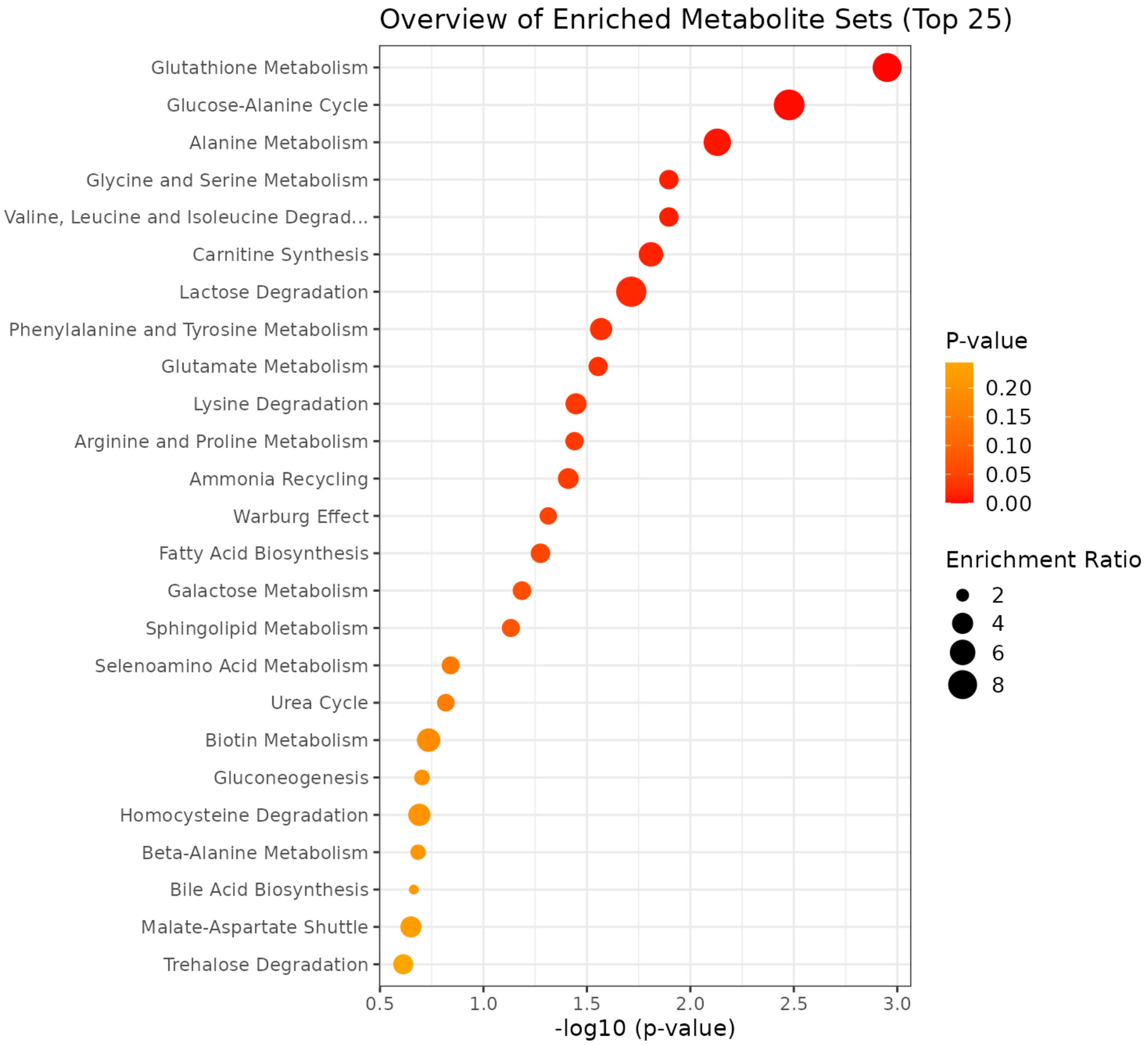

**Figure S10:** Pathway enrichment analysis of key metabolites altered under Trehalose treatment.

| Gene Name | Forward Primer (5'-3') | Reverse Primer (3'-5') |
|---|---|---|
| **GAPDH** | TGAGGCCGGTGCTGAGTATGTCG | GTGACGGTGGGTCTTCTGACACC |
| **VEGFA** | CTTGTTCAGAGCGGAGAAAGC | TTGCTTGCATGAACGTCTACA |
| **TWIST** | GTCCGCAGTCTTACGAGGAG | CTAAGTCTGGGAGTTCGACC |
| **ZEB1** | ACCTCTTCACAGGTTGCTCCT | ACTGAGAGTCGAGGACGTGA |
| **HIF-1α** | CTATGGAGGCCAGAAGAGGGTAT | AATACTCGGTGGACTACACCC |
| **VIMENTIN** | GACAATGCGTCTCTGGCACGTCTT | TTCTTGGACGTCCTCCGCCTCCT |
| **MCOX4** | TCTACTTCGGTGTGCCTTCG | CTTGTTCCCGTGGTTACTCA |

**Figure S11:** The figure depicts the list of Real Time PCR (qPCR) primers used in the study.

**Table S1:** The table depicting reagents, chemicals, commercial assay kits, public datasets, experimental models, oligonucleotides and software/algorithms used in the study.

| **Reagents** | **Source** | **Catalogue** |
|---|---|---|
| **Antibodies** | | |
| TFAM | Sigma-Aldrich | SAB1401383 |
| PGC-1α | Merck | ST1204 |
| CDK4 | Sigma-Aldrich | AV401 |
| p21 | Invitrogen | 337000 |
| Actin | | |
| Caspase-9 | Cell Signaling Technology | 52873T |
| Caspase-8 | Cell Signaling Technology | 9746L |
| Caspase-3 | Cell Signaling Technology | 9664T |
| Caspase-7 | Cell Signaling Technology | 12827T |
| PI3K | Cell Signaling Technology | 4292 |

| | | |
|---|---|---|
| Akt | Cell Signaling Technology | 2920S |
| p-Akt | Cell Signaling Technology | 4060L |
| p53 | Sigma-Aldrich | |
| Bcl-2 | Life Technologies | 138800 |
| B-catenin | Cell Signaling Technology | |
| AMPK | Sigma-Aldrich | SAB450233L |
| COX4 | | |
| c-Jun | Cell Signaling Technology | 9165L |
| MT-ND4 | Sigma-Aldrich | HPA053928 |
| HRP conjugated anti-rabbit | Sigma-Aldrich | |
| HRP conjugated anti-mouse | Sigma-Aldrich | |
| FITC-bound | Sigma-Aldrich | |
| Dy549-bound | Sigma-Aldrich | |
| AF488-bound | Sigma-Aldrich | |
| **Chemicals, peptides, and recombinant proteins** | | |
| Trehalose and Investigational Drugs | Sigma-Aldrich | |
| H2DCFDA | Molecular Probes | D399 |
| JC-1 Dye | | |
| Digitonin | Sigma-Aldrich | |
| Succinate | Sigma-Aldrich | |
| ADP | Sigma-Aldrich | 01905 |
| Cytochrome C | Sigma-Aldrich | |
| Oligomycin | Sigma-Aldrich | |
| FCCP | Sigma-Aldrich | |
| Rotenone | Sigma-Aldrich | |
| Antimycin A | Sigma-Aldrich | |
| Pyruvate | Sigma-Aldrich | |
| SYBR Green™ | Bio-Rad | |
| Fluoroshield mounting media containing DAPI | Thermofisher Scientific | |
| MitoTracker Green™ Probes | Thermofisher Scientific | M7514 |
| Ribitol | | |
| L-Norvaline | Sigma-Aldrich | |
| MSTFA | Sigma-Aldrich | |
| Western Blotting Chemicals | Sigma-Aldrich | |

| | | |
|---|---|---|
| PCR Chemicals | Thermofisher Scientific | |
| Protease Inhibitor Cocktail | | P8340 |
| Gibco$^{TM}$ DMEM (high glucose) | Invitrogen | 11995-073 |
| Fetal Bovine Serum | Invitrogen | F7524 |
| Pen-Strip Antibiotics | Invitrogen | 15140-122 |
| Trypsin (0.25%) | Invitrogen | 25200-056 |
| **Commercial assays Kits** | | |
| Cell Cycle Assay Kit | Thermofisher Scientific | BMS500PI |
| PI/Annexin V Assay | Thermofisher Scientific | A13199 |
| Live/Dead Cell Assay Kit | Thermofisher Scientific | |
| Revert-Aid cDNA Synthesis Kit | Thermofisher Scientific | K1622 |
| **Deposited data** | | |
| GEO2R Analyzed Data | This Paper | GSE215917; GSE174152; GSE68815; GSE219274; GSE223350; GSE234171; GSE188914 https://www.ncbi.nlm.nih.gov/geo/geo2r/ |
| GC-MS Data | This Paper | N/A |
| Compusyn Raw Data | This Paper | N/A |
| Combenefit Raw Data | This Paper | N/A |
| **Experimental models: Cell lines** | | |
| MCF7 | NCCS, Pune | N/A |
| MDA-MB-231 | NCCS, Pune | N/A |
| A549 | NCCS, Pune | N/A |
| T47D | NCCS, Pune | N/A |
| SHSHY5Y | NCCS, Pune | N/A |
| **Oligonucleotides** | | |
| Primers for ZEB1, Twist, Vimentin, VEGFA, Hif-1α | This Paper | Figure S11 |
| **Software and algorithms** | | |
| Combenefit Software | Combenefit [1] | https://sourceforge.net/projects/combenefit |
| Compusyn Software | Compusyn [2] | https://combosyn.com |
| C6 Plus Analyzer | BD Accuri | https://www.bdbiosciences.com |

| | | |
|---|---|---|
| Attune Cytometric Software version 7.1 | Thermofisher Scientific | https://www.thermofisher.com/in/en/home/life-science/cell-analysis/flow-cytometry.html |
| Datlab 8 | Oroboros Instruments | https://www.oroboros.at/index.php/product/datlab |
| GraphPad v8.0 | Prism | https://www.graphpad.com/features |
| Endnote v2025 | EndNote | https://endnote.com/downloads |
| FLUOVIEW FV1200/FV1000 Viewer | Olympus | https://olympus-fv1200-fv1000-viewer.updatestar.com/ |
| Image Lab Software | Bio-Rad | https://www.bio-rad.com/en-in/product/image-lab-software?ID=KRE6P5E8Z |
| CFX Maestro Software | Bio-Rad | https://www.bio-rad.com/en-in/product/cfx-maestro-software-for-cfx-real-time-pcr-instruments?ID=OKZP7E15 |
| **Other** | | |
